# Selective MOSPD2-STARD3 interaction at ER contact sites governs late endosome/lysosome dynamics and cholesterol homeostasis

**DOI:** 10.64898/2026.03.30.714413

**Authors:** Céline Knorr, Mehdi Zouiouich, Julie Eichler, Maxime Boutry, Corinne Wendling, Sophie Huver, Arthur Martinet, Catherine Fromental-Ramain, Elodie Monsellier, Guillaume Drin, Catherine Tomasetto, Fabien Alpy

**Author notes:** These authors contributed equally to this work (Céline Knorr, Mehdi Zouiouich). correspondence to Catherine Tomasetto, IGBMC, 1 rue Laurent Fries, 67400 Illkirch, France; and Fabien Alpy, IGBMC, 1 rue Laurent Fries, 67400 Illkirch, France.

## Abstract

Membrane contact sites (MCSs) between the endoplasmic reticulum (ER) and other organelles or the plasma membrane play essential roles in lipid exchange and organelle homeostasis; however, how the molecular composition of these sites is functionally specified remains poorly understood. Here, we identify the ER-resident protein MOSPD2 as a novel regulator of late endosome/lysosome (LE/Lys) homeostasis and show that MOSPD2 selectively recruits the cholesterol transporter STARD3 at ER-LE/Lys contacts. Loss of either MOSPD2 or STARD3 leads to an expansion of the LE/Lys compartment, accumulation of free cholesterol at LE/Lys, and impaired LE/Lys fusion dynamics, indicating that both proteins act within a common pathway at ER-LE/Lys contact sites.

STARD3 interacts with three members of the MSP-domain protein family, VAP-A, VAP-B and MOSPD2, yet, these tethers are not functionally equivalent at ER-LE/Lys MCSs. Using a series of quantitative binding assays, we show that STARD3 preferentially associates with MOSPD2 rather than with VAP-A or VAP-B, and that neither VAP-A nor VAP-B can compensate for the loss of MOSPD2 to restore LE/Lys biology and cholesterol distribution.

This work establishes the MOSPD2-STARD3 complex as a unique functional unit that regulates LE/Lys homeostasis. Collectively, these findings reveal that selective pairing between MSP-domain proteins and their FFAT-containing partners exist and shape the interaction landscape of MCSs to define their functions.

## Introduction

In eukaryotic cells, organelles are physically linked by membrane contact sites (MCSs) which are defined by the close juxtaposition of two organelle membranes, usually 10-30 nm apart, without fusion (Levine and Loewen, 2006; Scorrano et al., 2019). MCS are formed by tether proteins that bring membranes into close contact through protein-protein and/or protein-membrane interactions. This proximity between membranes facilitates the non-vesicular transfer of lipids and ions (Gatta and Levine, 2017; Wong et al., 2019; Prinz et al., 2020).

The endoplasmic reticulum (ER) is a dynamic, branched organelle that spreads throughout the cytosol and serves as a central hub for inter-organelle communication by establishing MCSs with virtually all other organelles. A small group of ER-resident proteins from the MSP (Major Sperm Protein) domain-containing family are key organizer of these contacts. These proteins known as VAP-A (Vesicle-Associated Membrane Protein-Associated Protein A), VAP-B, and MOSPD2 (MOtile SPerm Domain-containing protein 2), are anchored to the ER via a C-terminal transmembrane domain and extend their MSP domains toward the cytosol to recruit partner proteins associated to other organelles (Loewen et al., 2003; Di Mattia et al., 2018; Subra et al., 2023b; Levine, 2025). Through these physical associations, VAP-A, VAP-B or MOSPD2 drive the formation of MCSs between the ER and various organelles (Levine, 2025). Many of VAP-A, VAP-B and MOSPD2 protein partners carry an FFAT [two phenylalanines (FF) in an acidic tract (AT)] motif or a phospho-regulated Phospho-FFAT motif, that are selectively recognized by MSP domains (Loewen et al., 2003; Di Mattia et al., 2020a). The canonical FFAT motif follows the consensus sequence E_1_F_2_F_3_D_4_A_5_X_6_E_7_ (Mikitova and Levine, 2012) with two critical residues: the aromatic residue at position 2 that fits into a hydrophobic pocket of the MSP domain, and the acidic residue at position 4 that makes electrostatic contacts with basic residues of this domain (Kaiser et al., 2005; Di Mattia et al., 2020a; Furuita et al., 2010).

Proteomics databases suggest that VAP-A, VAP-B and MOSPD2 have hundreds of potential binding partners (Levine, 2025); however, the functional relevance of most of these interactions is unknown, with only a limited number of them being validated in cells. Given that VAP-A, VAP-B, and MOSPD2 are ubiquitously expressed and share many common partners, they may appear functionally redundant. Yet, genetic evidence strongly suggest that they are not fully interchangeable: VAP-A knockout (KO) mice die early during embryogenesis (McCune et al., 2017), whereas VAP-B– and MOSPD2 KO mice are viable (Kabashi et al., 2013; Yacov et al., 2020). These observations raise two major questions: Do members of the MSP protein family associate hierarchically with specific sets of partners? If so, does this hierarchy help organize the formation of diverse membrane contact sites with defined cellular functions?

One of the major organelles with which the ER establishes contacts is the late endosome and lysosome (LE/Lys), which serve as major nodes of the endocytic network, coordinating the sorting and degradation of endocytosed plasma membrane proteins and extracellular cargo, as well as intracellular material delivered through autophagy (Scott et al., 2014). LE/Lys form extensive contacts with the ER through multiple tethering complexes involving predominantly VAP-A, VAP-B and MOSPD2. Despite growing evidence that ER-LE/Lys contacts regulate processes such as cholesterol trafficking, organelle positioning, and signaling, how these contacts control LE/Lys biology remains poorly understood (Di Mattia et al., 2020b).

In this study, we investigated the functional redundancy among MSP-domain proteins in LE/Lys biology, and identified a specific role for MOSPD2. We show that loss of MOSPD2 leads to the expansion of the LE/Lys compartment through a mechanism that critically depends on its interaction with STARD3, a cholesterol transfer protein anchored at the LE/Lys membrane. We further demonstrate that the MOSPD2-STARD3 interaction defines functionally unique ER-LE/Lys contact sites that cannot be compensated for by STARD3 binding to VAP-A or VAP-B, and contributes to LE/Lys homeostasis. More globally, these findings reveal that selective MSP-domain tethers and their partners govern the organization and the functions of MCSs.

## Results

### MOSPD2 regulates late endosome/lysosome (LE/Lys) organization

To explore the role of MSP-domain proteins in LE/Lys biology, we individually silenced VAP-A, VAP-B, and MOSPD2, which are MSP-domain proteins with the known ability to bind FFAT-containing proteins. HeLa cells were treated with pools of siRNAs, and silencing efficiency was verified by Western blot (Fig. 1A). Cells were then labeled with the LE/Lys marker LAMP1 and imaged by immunofluorescence (Fig. 1B). In cells depleted of VAP-A or VAP-B, the number of LAMP1-positive vesicles was comparable to that in control cells (both untransfected and transfected with control siRNAs) (Fig. 1B, C). Strikingly, in cells silenced for MOSPD2 there was ∼2-fold more LAMP1-positive vesicles (Fig. 1B, C). To confirm this observation, we quantified LAMP1-positive vesicles in two independent clones of MOSPD2 knockout (KO) cells (MOSPD2 KO #1 and #2), which were generated using CRISPR/Cas9-mediated gene editing (Zouiouich et al., 2022). Again, in MOSPD2-deficient cells, the LAMP1-positive compartment was expanded by approximately twofold compared with control cells (Fig. 1B, C). Consistent with these data, LAMP1 protein levels were increased in MOSPD2-silenced and KO cells (Fig. 1A).

**Figure 1:**
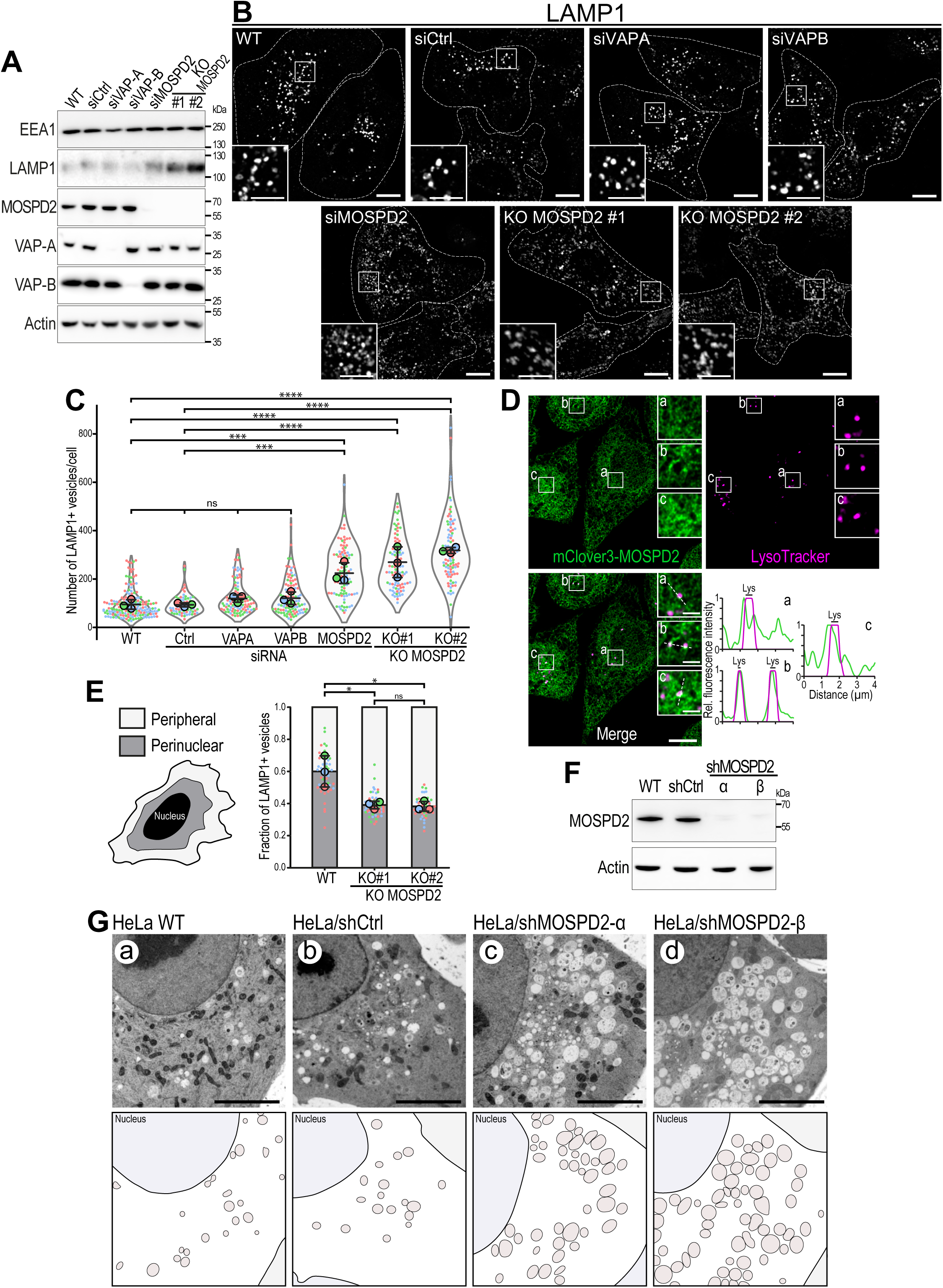
MOSPD2 regulates LE/Lys organization. A: Western blot analysis of EEA1 (early endosomes), LAMP1 (LE/Lys), MOSPD2 (ER), VAP-A (ER), VAP-B (ER) and actin protein levels in control HeLa cells (WT), HeLa cells transfected with control siRNAs (siCtrl), siRNAs targeting VAP-A (siVAP-A), VAP-B (siVAP-B), MOSPD2 (siMOSPD2), and in MOSPD2-deficient clones (KO#1 and KO#2). B: Representative confocal images of parental HeLa cells (WT), of HeLa cells transfected with control siRNAs (siCtrl), and with siRNAs targeting VAP-A (siVAP-A), VAP-B (siVAP-B), MOSPD2 (siMOSPD2), and MOSPD2-deficient clones (KO#1 and KO#2) labeled with anti-LAMP1 antibodies (LE/Lys, gray). Higher-magnification views of the areas outlined in white are shown. The contour of each cell is delimited by a white dotted line. Scale bars: 10 µm; inset scale bars: 5 µm. C: Quantification of LE/Lys number under conditions shown in B. Data are displayed as Superplots (Lord et al., 2020) showing the mean number of LE/Lys per cell (small dots) or per independent experiment (large dots). Independent experiments (n = 3) are color-coded. Means and error bars (SD) are shown as black bars. Data were collected from 181 (WT), 71 (siCtrl), 101 (siVAP-A), 109 (siVAP-B), 101 (siMOSPD2), 96 (KO#1), and 102 (KO#2) cells. One-way ANOVA with Tukey’s multiple comparisons test (ns, not significant; ***, P < 0.001; ****, P < 0.0001; n = 3 independent experiments). D: Live imaging of CRISPR/Cas9-edited HeLa cell expressing mClover3-MOSPD2 (green) at the endogenous levels and labelled with Lysotracker (magenta). Higher-magnification views of the areas outlined in white and labelled a-c are shown. Scale bar: 10 μm; inset scale bars: 2 µm. Lower right: line-scan analyses showing the relative fluorescence intensities of the magenta and green channels along the dotted lines in a-c. E: Quantification of LE/Lys spatial distribution in WT and MOSPD2-deficient HeLa cells labelled with anti-LAMP1 antibodies. LE/Lys vesicles were classified into two radial bins based on their normalized distance between the nuclear and cellular boundaries (left). Bar plots show the fraction of LE/Lys per bin, displaying the mean perinuclear fraction per cell (small dots) and per independent experiment (large dots). Independent experiments (n=3) are color-coded. Means and SD are indicated as black bars. Data were collected from 77 (WT), 76 (KO #1), and 75 (KO #2) cells. One-way ANOVA with Tukey’s multiple comparisons test (ns, not significant; *, P < 0.05; n = 3 independent experiments). F: Western blot analysis of MOSPD2 protein levels in control HeLa cells (WT), HeLa cells transfected with control shRNAs (shCtrl) and two individual shRNAs targeting MOSPD2 (shMOSPD2-α and –β). Actin was used as a loading control. G: TEM images of control HeLa cells (a: HeLa; b: HeLa/shCtrl) and MOSPD2-silenced HeLa cells (c: HeLa/sh-MOSPD2-α; d: HeLa/shMOSPD2-β). An interpretation scheme is shown on the bottom; endosomes are in light gray. Scale bars: 5 μm.

MOSPD2 is an ER-resident protein involved in scaffolding diverse ER-organelle contact sites. Our previous data suggest a preferential involvement in the formation of ER-LE/Lys and ER-LD contacts (Di Mattia et al., 2018; Zouiouich et al., 2022). To further investigate the distribution of MOSPD2 within the cell and its role in organelle interactions, we imaged HeLa cells in which the endogenous MOSPD2 gene was modified to encode an in-frame fusion with the fluorescent protein mClover3 (Zouiouich et al., 2022). Consistent with previous findings, mClover3-MOSPD2 displayed a distinctive reticular pattern throughout the cytoplasm, typical of ER morphology (Fig. 1D). Additionally, we observed that MOSPD2 was enriched in discrete foci that co-localized with Lysotracker, a fluorescent marker that selectively labels acidic endosomal compartments, i.e. LE/Lys (Fig. 1D). This suggests that a significant fraction of MOSPD2 is localized at ER-LE/Lys contact sites and may play a specific role in LE/Lys biology. In addition to LAMP1, LE/Lys are characterized by the presence of a unique lipid species, bis(monoacylglycero)phosphate (BMP), a low intraluminal pH, and high degradation activity (Gruenberg, 2001). Thus, we analyzed whether the amplified endocytic structures seen in MOSPD2-deficient cells possess these specific LE/Lys features (Fig. S1). In cells lacking MOSPD2, the number of vesicles labeled with i) antibodies against BMP (Fig. S1A), ii) Lysotracker (Fig. S1B), and iii) DQ-Red BSA, a fluorogenic substrate reflecting lysosomal protease activity (Fig. S1C), was significantly increased. We then wondered whether the early endocytic compartment was also affected by the loss of MOSPD2 expression. We labeled control (WT) and MOSPD2 KO cells with antibodies against the early endosome antigen 1 (EEA1) and found that the number of early endosomes was not modified in cells devoid of MOSPD2 compared to control cells (Fig. S1D).

We then investigated whether the spatial distribution of LE/Lys was altered in MOSPD2-deficient cells (Fig. 1E). To this end, cells were segmented based on the contours of the nucleus and the cell, and the cytoplasm was divided into two regions: a perinuclear region and a peripheral region. The fraction of LE/Lys located in each region was quantified. In control cells, LE/Lys were mainly perinuclear, whereas in MOSPD2-deficient cells they were redistributed toward the cell periphery, indicating that loss of MOSPD2 alters the sub-cellular distribution of LE/Lys.

We further examined the phenotype of MOSPD2-deficient cells by an ultrastructural approach; HeLa cells were silenced for MOSPD2 using small hairpin RNAs (shRNAs) (Fig. 1G) and then processed for transmission electron microscopy (TEM). We observed an increase in the number of multivesicular vesicles corroborating the immunofluorescence analysis.

To assess whether this phenotype was specific to HeLa cells or reflected a broader role of MOSPD2 in endosome regulation, we quantified LAMP1-positive structures in the immortalized fibroblast cell line MRC5 following MOSPD2 silencing (Fig. S2A). As in HeLa cells, MOSPD2 depletion led to an increased number of LAMP1-positive structures, suggesting that the expansion of the LE/Lys compartment upon MOSPD2 loss is a general phenomenon.

Altogether, these data show that among proteins from the VAP family, MOSPD2 has a non-redundant function on LE/Lys since only cells lacking MOSPD2 exhibit an expansion of the LE/Lys compartment, supporting the notion that the ER-anchored protein MOSPD2 contributes to the homeostasis of these organelles.

### MOSPD2-deficient cells exhibit cholesterol-enriched LE/Lys

As LE/Lys carry out key metabolic roles in cells, notably in cholesterol homeostasis (Ikonen and Olkkonen, 2023), and since these functions are linked to their intracellular positioning (Cabukusta and Neefjes, 2018), we wondered whether the LE/Lys phenotype observed in MOSPD2 KO cells was associated with changes in cholesterol distribution. To address this, we labeled free cholesterol using the fluorescent probe filipin (Wüstner et al., 2012) (Fig. 2A-C). In control cells, free cholesterol was predominantly localized at the plasma membrane, as expected (Eichler et al., 2025), with weaker labeling of a limited number of intracellular compartments, most likely the Golgi apparatus and endosomes (Fig. 2A). In MOSPD2-deficient cells, in addition to plasma membrane staining, we observed numerous intracellular structures displaying strong filipin staining (Fig. 2B, C). Co-labeling with LAMP1 showed that many of these filipin-positive intracellular structures were LE/Lys (Fig. 2A-C). Moreover, quantification of filipin labeling in LAMP1-positive vesicles (Fig. 2D) revealed higher intensity in MOSPD2-deficient cells relative to control cells, suggesting that free cholesterol is enriched in LE/Lys when MOSPD2 is absent.

**Figure 2:**
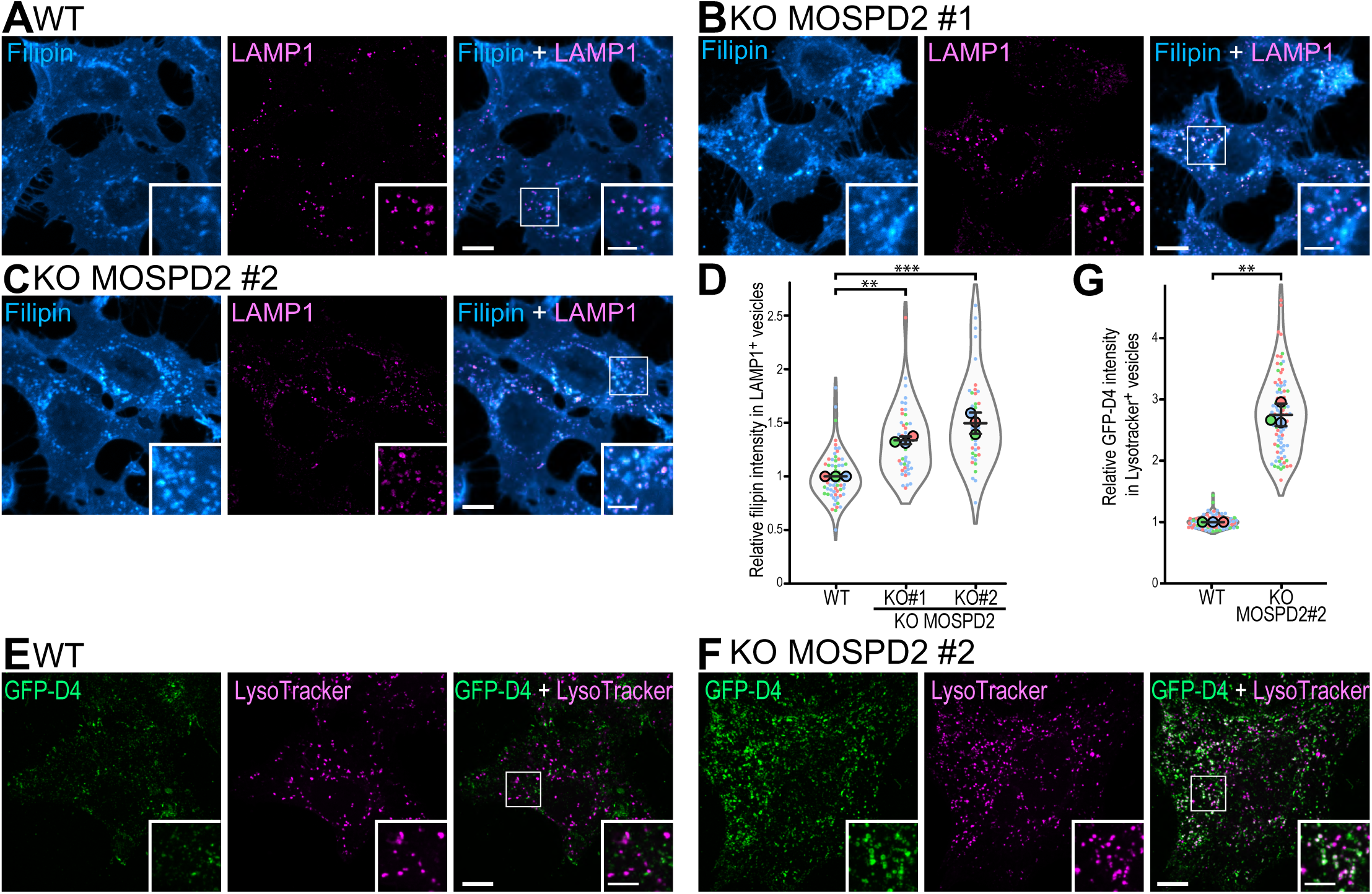
MOSPD2 regulates intracellular cholesterol localization. A-C: WT (A) and MOSPD2-deficient (KO MOSPD2 #1 in B; KO MOSPD2 #2 in C) HeLa cells were labeled with anti-LAMP1 antibodies (magenta) and the fluorescent cholesterol probe filipin (Cyan Hot). Higher magnifications images of the area outlined in white are shown on the right. Scale bars: 10 µm; inset scale bars: 5 µm. D: Quantification of relative filipin fluorescence intensity in LAMP1-positive vesicles for the conditions shown in A-C. Data are presented as Superplots, displaying the mean number of LE/Lys per cell (small dots) and per independent experiment (large dots). Independent experiments (n = 3) are color-coded. Means and SD are indicated as black bars. Data were collected from 70 (WT), 46 (KO MOSPD2 #1), and 42 (KO MOSPD2 #2) cells. One-way ANOVA with Tukey’s multiple comparisons test (**, P < 0.01; ***, P < 0.001; n = 3 independent experiments). E, F: WT (E) and MOSPD2-deficient (F) HeLa cells were labeled with the fluorescent cholesterol probe GFP-D4 (green), and Lysotracker (magenta). Higher magnifications images of the area outlined in white are shown. Scale bars: 10 µm; inset scale bars: 5 µm. G: Quantification of relative GFP-D4 fluorescence intensity in Lysotracker-positive vesicles for the conditions shown in E-F. Data are presented as Superplots, displaying the normalized mean GFP-D4 fluorescence intensity per cell (small dots) and per independent experiment (large dots). Independent experiments (n = 3) are color-coded. Means and SD are indicated as black bars. Data were collected from 97 (WT), and 89 (KO MOSPD2 #2) cells. Student’s unpaired t-test (**, P < 0.01; n = 3 independent experiments).

To confirm this observation, we used another cholesterol probe consisting of the D4 domain of perfringolysin O (θ toxin, *Clostridium perfringens*) fused to EGFP (Ohno-Iwashita et al., 2004; Wilhelm et al., 2019). We labelled intracellular structures with the EGFP-D4 probe in permeabilized cells (Fig. 2E-G). In control cells, only a few vesicles were positive for EGFP-D4 labeling, some of which were Lysotracker-positive (Fig. 2E). By contrast, MOSPD2-deficient cells displayed many EGFP-D4– and Lysotracker-positive vesicles (Fig. 2F, G) further supporting increased cholesterol levels in LE/Lys in MOSPD2-deficient cells.

Together, these data indicate that MOSPD2 is required for the normal distribution of free cholesterol, and that when the protein is missing, cholesterol is enriched at LE/Lys.

### MOSPD2 functions on LE/Lys via its FFAT-binding MSP domain

Having established that expression of the ER-resident MOSPD2 protein impacts LE/Lys number, positioning and sterol content, we asked whether it does so via its presence at ER-LE/Lys contacts, or via a more indirect process. To determine which activity of MOSPD2 is involved in LE/Lys biology, we performed rescue experiments by restoring MOSPD2 expression in KO cells (MOSPD2 KO#2) using mScarlet-tagged WT or mutant MOSPD2 (Fig. 3A, B). In line with data presented in Figure 1, cells lacking MOSPD2 had 2 times more LE/Lys compared to WT cells (Fig. 3Ca, b and D). When mScarlet-MOSPD2 was re-expressed in KO cells, the number of LE/Lys was similar to that in WT cells (Fig. 3Cc and D). Within the MSP domain-containing family, MOSPD2 stands out in that, in addition to the MSP domain, it contains a CRAL-TRIO [cellular retinaldehyde-binding protein (CRALBP) and triple functional domain protein (TRIO)] domain, which is specifically involved in ER-lipid droplet (LD) contact formation and LD homeostasis (Zouiouich et al., 2022). Expression of MOSPD2 mutants either devoid of the CRAL-TRIO domain (mScarlet-MOSPD2 ΔCRAL-TRIO) or with a CRAL-TRIO domain unable to associate with lipid droplets (mScarlet-MOSPD2 W201E) also rescued the KO phenotype (Fig. 3Cd, e and D), indicating that the CRAL-TRIO domain is dispensable for MOSPD2 function at LE/Lys. In contrast, expression of MOSPD2 mutants either lacking the MSP domain (mScarlet-MOSPD2 ΔMSP), or with the RD/LD mutation, defective in recognizing FFAT motifs (mScarlet-MOSPD2 RD/LD) (Di Mattia et al., 2018), failed to rescue the abnormal LE/Lys phenotype seen in MOSPD2 KO cells (Fig. 3Cf, g and D).

**Figure 3:**
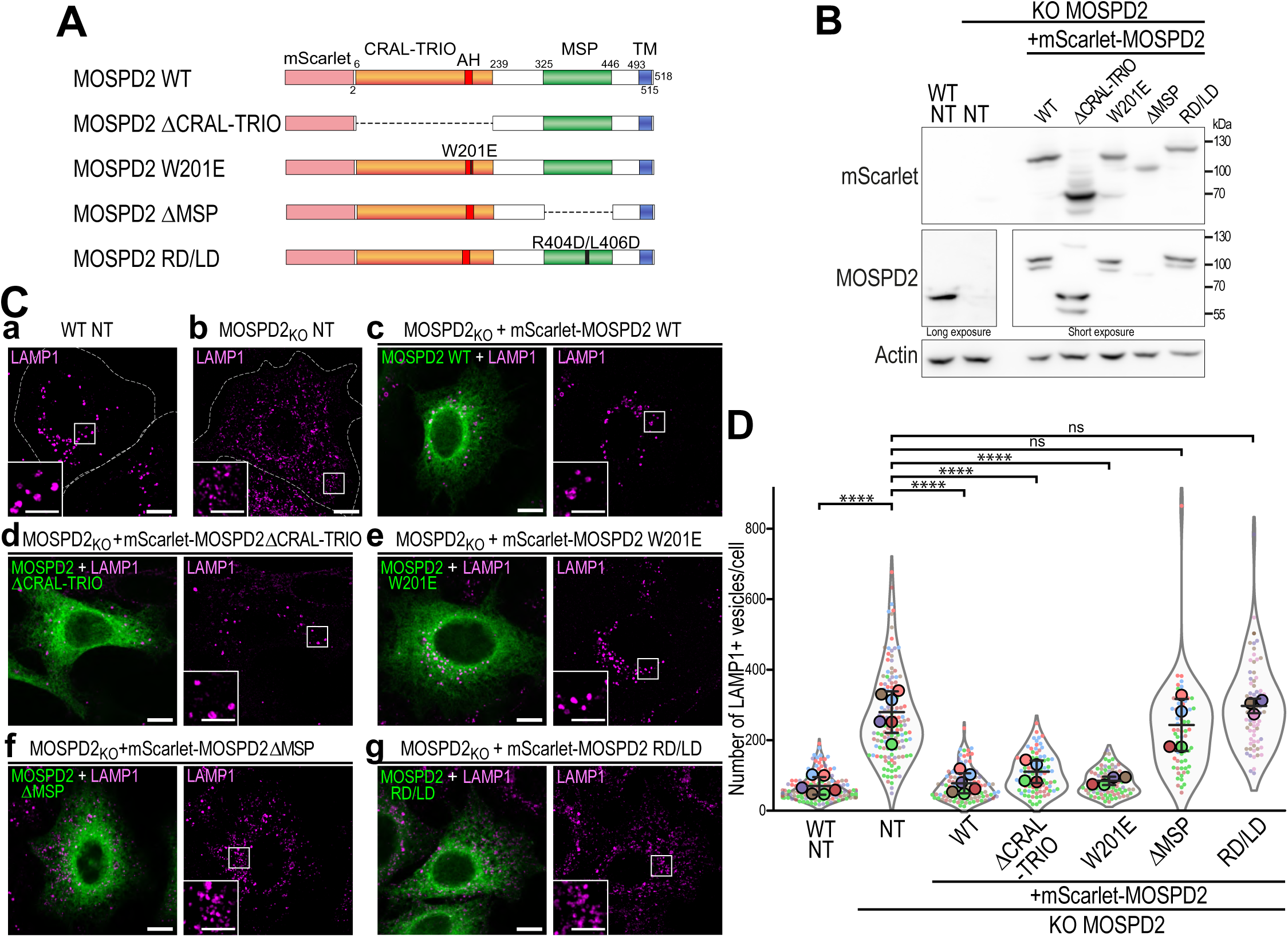
The MSP domain of MOSPD2 is required for proper LE/Lys organization. A: Schematic representation of the different WT and mutant proteins used in the study, including domain deletions (ΔCRAL-TRIO, ΔMSP) and point mutations (W201E, RD/LD) impairing the CRAL-TRIO and MSP domain functions, respectively. B: Western Blot analysis of mScarlet, MOSPD2 and actin protein levels in WT and MOSPD2 KO HeLa cells. MOSPD2 expression was rescued in MOSPD2 KO cells using mScarlet-MOSPD2 expression constructs shown in A. NT, non-transfected. Endogenous MOSPD2 expression is shown after a long exposure. C: Representative confocal images of non-transfected (NT) WT (a) and MOSPD2 KO (b) HeLa cells, and of MOSPD2 KO cells in which MOSPD2 expression was restored using the mScarlet-MOSPD2 constructs (c-g; green) depicted in panel A. LE/Lys were stained with anti-LAMP1 antibodies (magenta). Higher magnifications images of the area outlined in white are shown. Scale bars: 10 µm; inset scale bars: 5 µm. D: Quantification of LE/Lys numbers in cells shown in C. Data are displayed as Superplots showing the mean number of LAMP1-postive LE/Lys per cell (small dots) or per independent experiment (large dots). Independent experiments (n = 3-6) are color-coded. Means and error bars (SD) are shown as black bars. Data were collected from 153 (WT), 147 (KO MOSPD2), 112 (KO transfected by MOSPD2), 91 (KO transfected by MOSPD2 ΔCRAL-TRIO), 85 (KO transfected by MOSPD2 W201E), 69 (KO transfected by MOSPD2 ΔMSP), and 74 (KO transfected by MOSPD2 RD/LD) cells. One-way ANOVA with Tukey’s multiple comparisons test (ns, not significant; ****, P < 0.0001; n = 3-6 independent experiments).

Since the MSP domain mediates interactions with FFAT-containing proteins and the formation of ER-organelle contacts, these data suggested that MOSPD2 acts at ER-LE/Lys contacts to maintain lysosomal homeostasis.

### STARD3 loss phenocopies MOSPD2 deficiency

The finding that MOSPD2 acts on LE/Lys homeostasis through its MSP domain suggests that it recruits a specific LE/Lys protein that, either directly and/or through the formation of ER-LE/Lys MCS, modulates LE/Lys biology. Furthermore, as MOSPD2 deficiency leads to abnormal cholesterol enrichment in LE/Lys, we hypothesized that such a partner might be involved in cholesterol homeostasis. We therefore focused on three known cholesterol transfer proteins localized to LE/Lys known to interact with MOSPD2: two members of the ORD [oxysterol-binding protein(OSBP)-related domain] LTP family: OSBP (Olkkonen, 2015; Mesmin et al., 2013; Lim et al., 2019; Cabukusta et al., 2020) and OSBP-related protein 1 (ORP1) (Johansson et al., 2005; Rocha et al., 2009; Johansson et al., 2002; Di Mattia et al., 2018), and STARD3 [StAR-related lipid transfer (START) domain protein 3, also known as metastatic lymph node 64, MLN64], a member of the START LTP family (Alpy et al., 2013; Wilhelm et al., 2017; Voilquin et al., 2019; Di Mattia et al., 2020a) (Fig. 4A). To determine if ORP1L, OSBP or STARD3 work in concert with MOSPD2 to regulate LE/Lys homeostasis, we silenced these genes individually in HeLa cells (Fig. 4B) and labeled LE/Lys with anti-LAMP1 antibodies (Fig. 4C).

**Figure 4:**
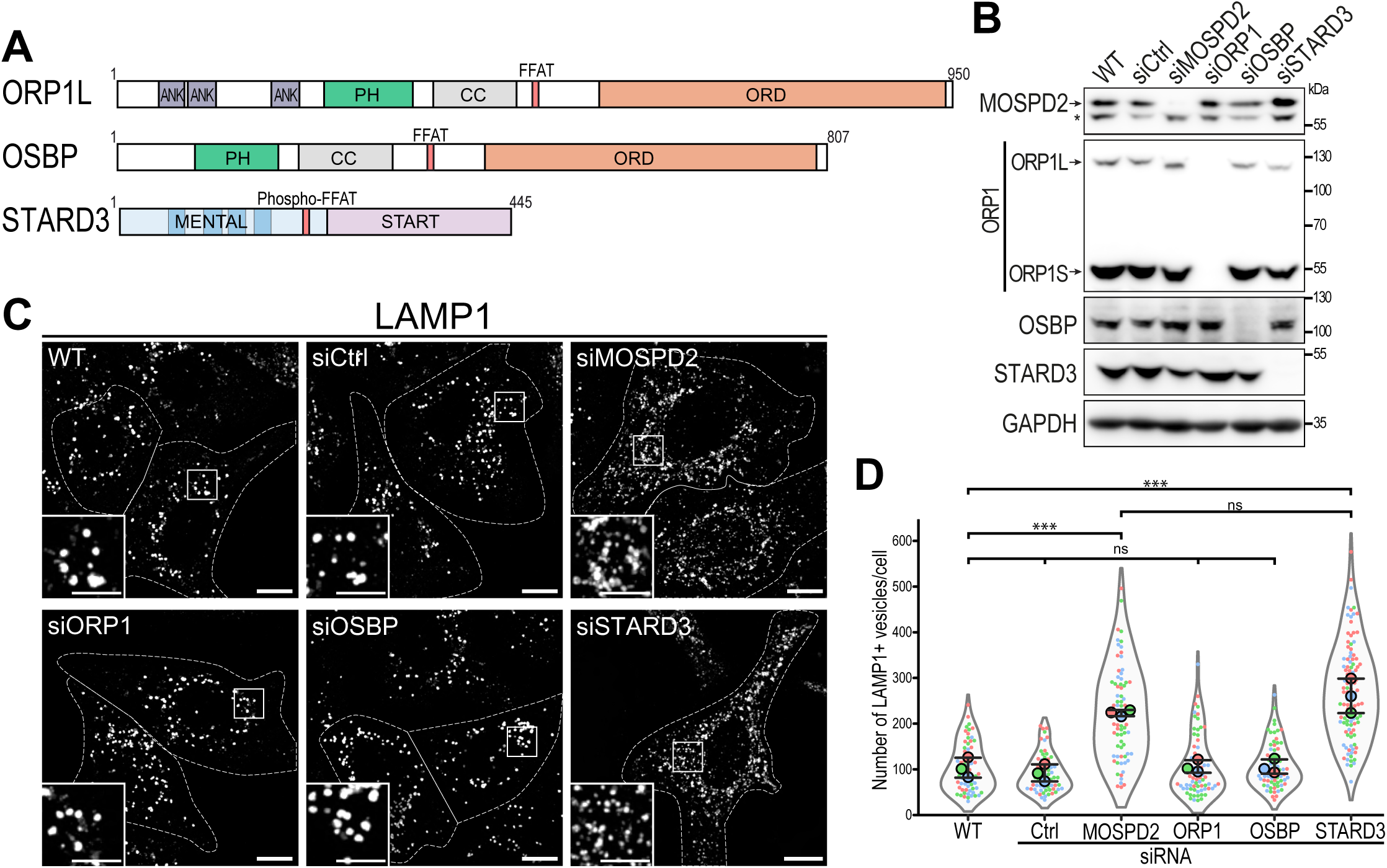
Targeted screening identifies STARD3 as a MOSPD2 partner whose loss phenocopies MOSPD2 deficiency. A: Primary structure of ORP1L, OSBP, and STARD3. ANK: ankyrin repeats; Pleckstrin Homology (PH) domain; ORD: OSBP-related ligand-binding domain; CC: coiled-coil; START: StAR-related lipid-transfer domain; MENTAL: MLN64 NH_2_-terminal domain. B: Western Blot analysis of MOSPD2 (*, aspecific), ORP1 (ORP1L, long isoform; ORP1S short isoform), OSBP, STARD3 and GAPDH in control HeLa cells (WT and siCtrl) and in cells treated with siRNAs targeting MOSPD2, ORP1, OSBP, and STARD3. C: Confocal images of LAMP1 labeling in control HeLa cells (WT and siCtrl) and in cells treated with siRNAs targeting MOSPD2, ORP1, OSBP and STARD3. The contour of each cell is delimited by a white dotted line. Higher magnifications images of the area outlined in white are shown. Scale bars: 10 µm; inset scale bars: 5 µm. D: Quantification of LE/Lys numbers under the conditions shown in panel C. Data are displayed as Superplots showing the mean number of LE/Lys per cell (small dots) and per independent experiment (large dots). Independent experiments (n = 3) are color-coded. Means and error bars (SD) are shown as black bars. Data were collected from 67 (WT), 66 (siCtrl), 70 (siMOSPD2), 72 (siORP1), 73 (siOSBP), and 109 (siSTARD3) cells. One-way ANOVA with Tukey’s multiple comparisons test (ns, not significant; ***, P < 0.001; n = 3 independent experiments).

Silencing ORP1 or OSBP did not modify the number of LAMP1-positive vesicles. In contrast, STARD3 depletion led to a ∼2-fold increase in LE/Lys compared to WT cells (Fig. 4C, D), mirroring the phenotype observed with MOSPD2 silencing (Fig. 4C, D).

To confirm the impact of STARD3 on LE/Lys homeostasis, we inactivated STARD3 in HeLa cells using CRISPR/Cas9-mediated gene editing. Two independent KO clones (STARD3 KO#1 and #2) were analyzed (Fig. 5A). Consistent with the siRNA results, we found that these cells exhibited a marked increase in the number of LAMP1-positive vesicles (Fig. 5B, C), which were more present in the cell periphery (Fig. 5D). This was accompanied by increased LAMP1 and LAMP2 protein levels in STARD3 KO cells (Fig. 5A). Similarly, the number of Lysotracker-positive vesicles was increased in STARD3 KO cells (Fig. S2A).

**Figure 5:**
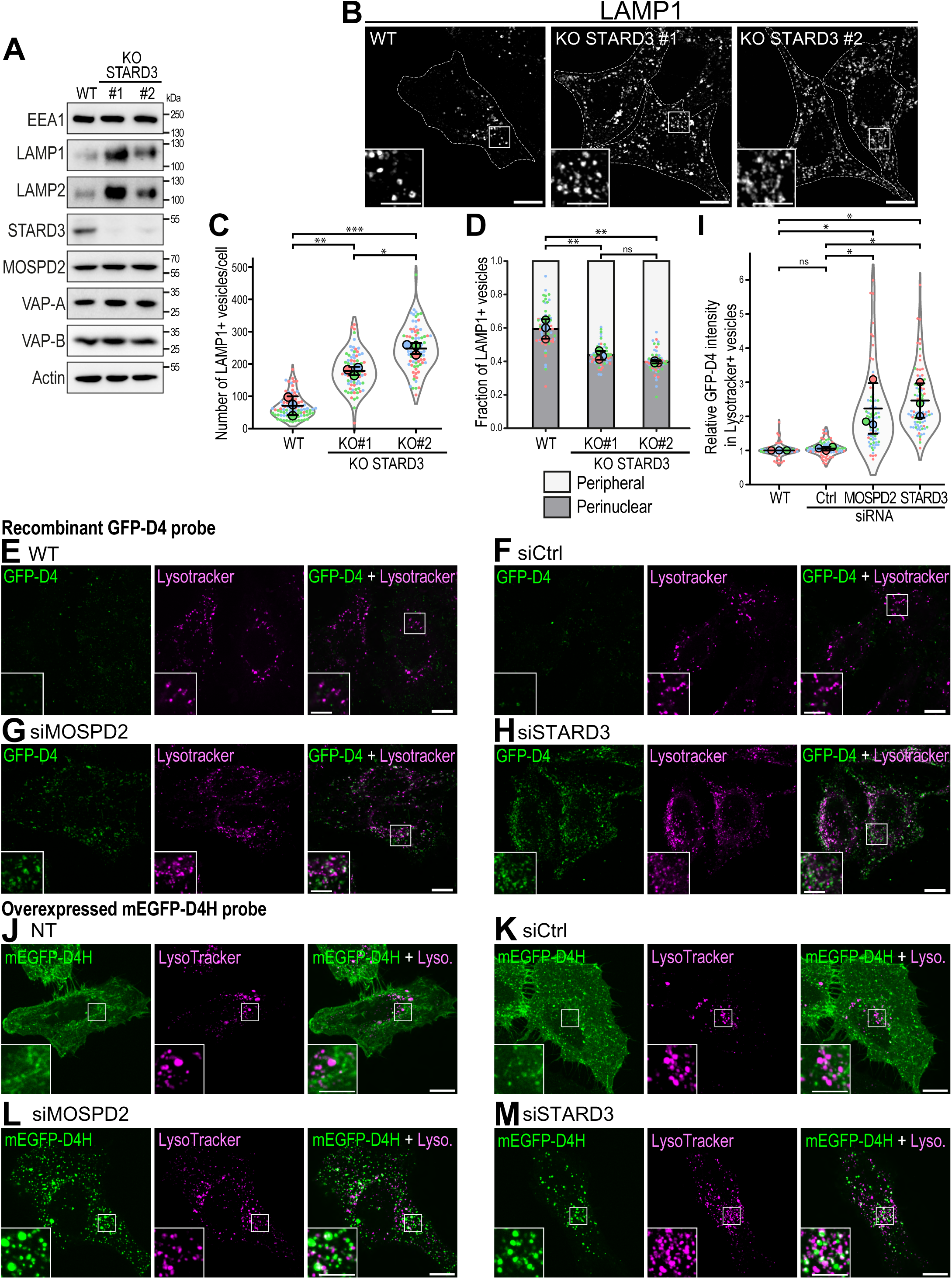
STARD3 loss phenocopies MOSPD2 deficiency. A: Western Blot analysis of EEA1, LAMP1, LAMP2, STARD3, MOSPD2, VAP-A, VAP-B and Actin in HeLa cells WT and in STARD3-deficient clones (#1 and #2). B: Confocal images of LAMP1 labeling in WT HeLa cells and STARD3-deficient cells (KO STARD3 #1 and KO STARD3 #2). The contour of each cell is delimited by a white dotted line. Higher magnifications images of the area outlined in white are shown. Scale bars: 10 µm; inset scale bars: 5 µm. C: Quantification of LE/Lys numbers under the conditions shown in panel B. Data are displayed as Superplots showing the mean number of LE/Lys per cell (small dots) and per independent experiment (large dots). Independent experiments (n = 3) are color-coded. Means and error bars (SD) are shown as black bars. Data were collected from 92 (WT), 80 (KO STARD3 #1), and 83 (KO STARD3 #2) cells. One-way ANOVA with Tukey’s multiple comparisons test (*, P < 0.05; **, P < 0.01; ***, P < 0.001; n = 3 independent experiments). D: Quantification of LE/Lys spatial distribution in WT and STARD3-deficient HeLa cells labelled with anti-LAMP1 antibodies. Bar plots show the fraction of LE/Lys per bin, displaying the mean perinuclear fraction per cell (small dots) and per independent experiment (large dots). Independent experiments (n=3) are color-coded. Means and SD are indicated as black bars. Data were collected from 94 (WT), 78 (KO #1), and 82 (KO #2) cells. One-way ANOVA with Tukey’s multiple comparisons test (ns, not significant; **, P < 0.01; n = 3 independent experiments). E-H: HeLa cells either untransfected (E) or transfected with control siRNAs (F), siRNAs targeting MOSPD2 (G), or STARD3 (H) were labeled with the fluorescent cholesterol probe GFP-D4 (green), and Lysotracker (magenta). Higher magnifications images of the area outlined in white are shown. Scale bars: 10 µm; inset scale bars: 5 µm. I: Quantification of relative GFP-D4 fluorescence intensity in Lysotracker-positive vesicles under the conditions shown in E-H. Data are presented as Superplots, displaying the mean GFP-D4 fluorescence intensity per cell (small dots) and per independent experiment (large dots). Independent experiments (n = 3) are color-coded. Means and SD are indicated as black bars. Data were collected from 79 (WT), 79 (siCtrl), 81 (siMOSPD2), 95 (siSTARD3) cells. One-way ANOVA with Tukey’s multiple comparisons test (*, P < 0.05; n = 3 independent experiments). J-M: HeLa cells stably expressing the fluorescent cholesterol probe mEGFP-D4H (green), either untransfected (J) or transfected with control siRNAs (K), siRNAs targeting MOSPD2 (L), or STARD3 (M) were labeled with Lysotracker (magenta). Higher magnification images of the area outlined in white are shown. Scale bars: 10 µm; inset scale bars: 5 µm.

Given that MOSPD2 KO cells display cholesterol accumulation in LE/Lys, we investigated whether loss of STARD3 would produce a similar effect. Cholesterol was labeled with filipin in cells treated with siRNAs targeting MOSPD2 and STARD3. As negative control, cells were treated with control siRNA and siRNA targeting VAP-A. In agreement with Figure 2, MOSPD2 silencing led to increased filipin labeling in LAMP1-positive LE/Lys (Fig. S3 A-C and F). A similar phenotype was observed with STARD3 silencing, which resulted in cholesterol buildup in LAMP1-positive LE/Lys (Fig. S3 D, F). In contrast, VAP-A silencing had no effect on cholesterol localization (Fig. S3 E, F). These results were further validated by intracellular cholesterol labeling using the GFP-D4 probe (Fig. 5E-I). Cells were stained with lysotracker, then fixed, permeabilized, and incubated with the GFP-D4 probe. Compared with control cells (Fig. 5E, F, I), cells silenced for MOSPD2 (Fig. 5G, I) or STARD3 (Fig. 5H, I) displayed numerous LE/Lys positive for the GFP-D4 probe. These findings were confirmed in STARD3 KO cells, which also exhibited increased intracellular GFP-D4H staining (Fig. S3 G-H). To determine whether cholesterol accumulates at the limiting membrane of LE/Lys or within the lumen, we stably expressed a variant of the D4 probe named D4H fused to monomeric EGFP in HeLa cells. In control cells (Fig. 5I, J), the mEGFP-D4H probe predominantly labeled the plasma membrane, with weaker labeling of intracellular structures. In contrast, in MOSPD2– or STARD3-silenced cells (Fig. 5K, L and Fig. S3I), the probe strongly labeled intracellular vesicles, often Lysotracker-positive, while the plasma membrane staining was markedly reduced, indicating that cholesterol accumulates at the limiting membrane of LE/Lys in MOSPD2– and STARD3-silenced cells. Together, these results show that both MOSPD2 and STARD3 regulate cholesterol levels in LE/Lys.

Thus, the loss of STARD3, a LE/Lys partner of MOSPD2, mimics the LE/Lys phenotype observed in the absence of MOSPD2, supporting the idea that MOSPD2 and STARD3 function together to regulate the biology of LE/Lys and cholesterol trafficking.

### MOSPD2– and STARD3-deficient cells display LE/Lys fusion defects

The dynamics of the endocytic pathway rely on a balance between membrane fusion and fission events involving tubules and vesicles (Yang and Wang, 2021). The increased number of LE/Lys observed in MOSPD2– and STARD3-deficient cells could therefore result from increased fission, reduced fusion, or a combination of both.

To determine whether LE/Lys fission is altered upon MOSPD2 depletion, we monitored fission events in living cells expressing mCherry-LAMP1. Tubule formation followed by fission is a major mode of LE/Lys remodeling and reformation (Yang and Wang, 2021). U2OS cells were transfected with either control siRNAs or siRNAs targeting MOSPD2 and subjected to amino acid starvation for 10 h in Hanks’ Balanced Salt Solution (HBSS) to stimulate the formation and fission of LE/Lys tubules, as previously described (Boutry et al., 2023). Live-cell imaging was then performed, and the number of mCherry-LAMP1-positive tubules undergoing fission was quantified (Fig. 6A). The proportion of fission events was comparable between control and MOSPD2-silenced cells (Fig. 6A), indicating that LE/Lys tubule fission rates were not significantly affected by MOSPD2 loss.

**Figure 6:**
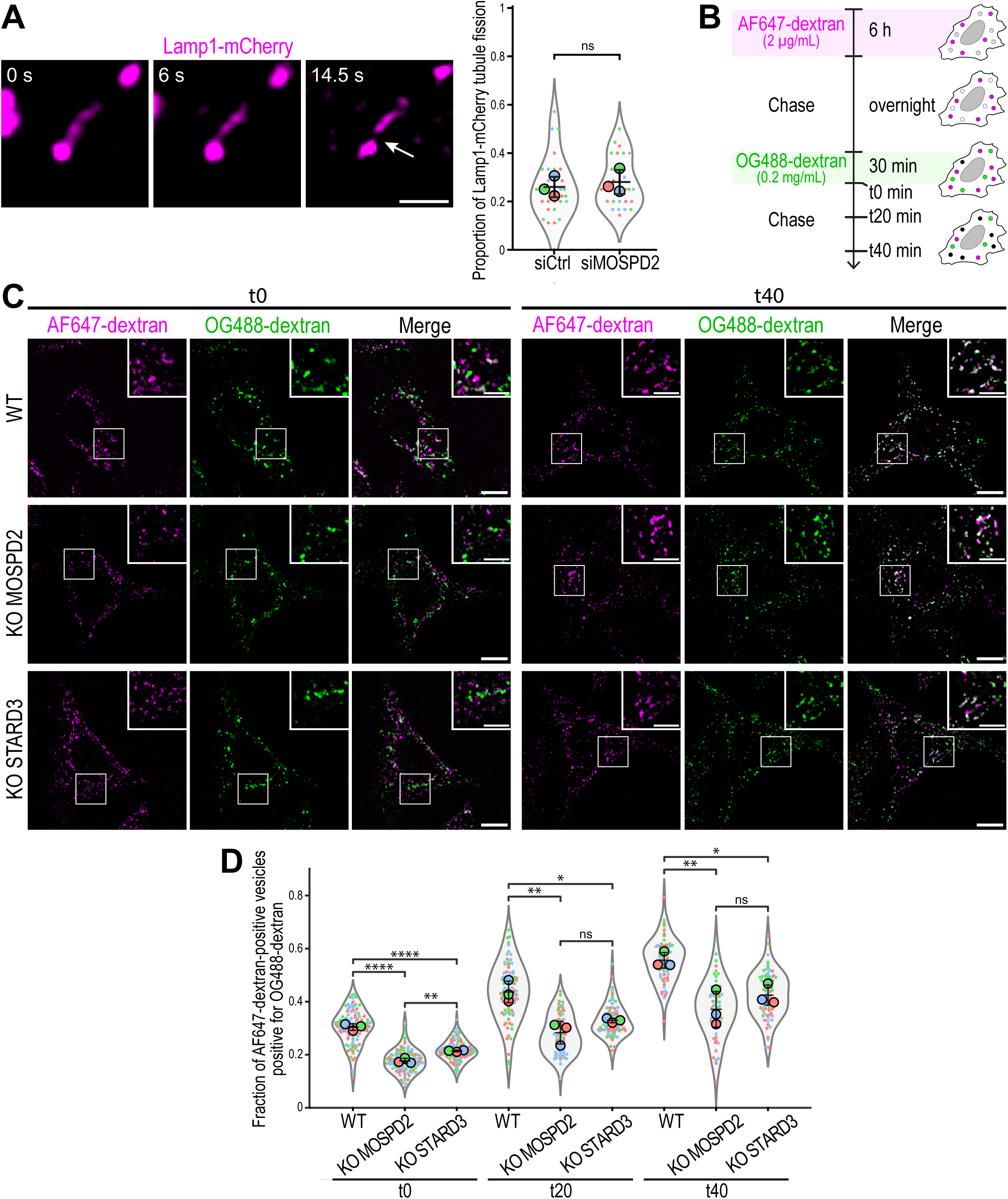
MOSPD2– and STARD3-deficient cells display LE/Lys fusion defects. A: Left: illustration of a fission event (arrow) in U2OS cells expressing Lamp1-mCherry. Scale bar: 1 µm. Right: quantification of fission events in control HeLa cells (siCtrl) and in cells treated with siRNAs targeting MOSPD2 (siMOSPD2). Data are presented as Superplots. displaying the proportion of Lamp1-mCherry tubule undergoing fission per cell (small dots) and per independent experiment (large dots). Independent experiments (n = 3) are color-coded. Means and SD are indicated as black bars. Data were collected from 28 (siCtrl), and 25 (siMOSPD2) cells. Student’s unpaired t-test (ns, not significant; n = 3 independent experiments). B: Principle of the dual-color dextran pulse-chase assay. Cells were sequentially incubated with fluorescent dextrans: first, far-red dextran (AF647-dextran) for several hours. After an overnight chase to label LE/Lys, cells were pulse labelled for 30 min with a green dextran (OG488-dextran) to label early endocytic compartments destined to fuse with LE/Lys. Co-localization between far-red and green dextrans within endocytic vesicles was quantified either immediately after the second pulse (t0) or after 20– or 40-min chase periods to monitor the progressive fusion of endocytic compartments over time. C: Confocal images of WT, MOSPD2-deficient, and STARD3-deficient HeLa cells incubated with Alexa Fluor 647-labeled dextran (AF647-dextran; magenta), and Oregon Green-labeled dextran (OG488-dextran; green), either without chase (t0) or after a 40-min chase (t40). Scale bar: 10 µm; inset scale bars: 5 µm. D: Quantification of the fraction of AF647-dextran-positive vesicles positive for OG488-dextran without chase (t0) or after a 20– (t20) or a 40-min chase (t40). Data are displayed as Superplots showing the fraction per cell (small dots) and per independent experiment (large dots). Independent experiments (n = 3) are color-coded. Means and error bars (SD) are shown as black bars. Data were collected from 94 (WT t0), 76 (WT t20), 68 (WT t40), 89 (KO MOSPD2 t0), 68 (KO MOSPD2 t20), 62 (KO MOSPD2 t40), 89 (KO STARD3 t0), 78 (KO STARD3 t20), and 75 (KO STARD3 t40) cells. One-way ANOVA with Tukey’s multiple comparisons test (ns; not significant; *, P < 0.05; **, P < 0.01; ****, P < 0.0001; n = 3 independent experiments).

We next assessed whether LE/Lys fusion was impaired in MOSPD2-deficient cells. To do so, we used a dual-color dextran pulse-chase assay. Briefly, sequential uptake of fluorescent dextrans, which enter cells via endocytosis, allowed us to distinguish pre-existing late endocytic/lysosomal compartments (AF647-dextran positive) from early endocytic vesicles (OG488-dextran positive) (Fig. 6B). Co-localization between far-red and green dextrans within endocytic vesicles was quantified either immediately after the second pulse (t0) or after 20– or 40-min chase periods to monitor the progressive fusion of endocytic compartments over time (Fig. 6C). Compared with WT cells, a reduced colocalization was observed in MOSPD2-deficient cells at all time points analyzed (t0, t20 and t40) (Fig. 6D). Although colocalization increased over time in both conditions, it remained consistently lower in MOSPD2-deficient cells, indicating that the rate of endosomal fusion was reduced.

We performed the same experiment in STARD3-deficient cells (Fig. 6B, C). Consistent with previous reports showing impaired LE/Lys fusion upon STARD3 loss (Holtta-Vuori et al., 2005), STARD3-deficient cells displayed fusion defects similar to those observed in MOSPD2-deficient cells.

Together, these results suggest that reduced LE/Lys fusion underlies the expansion of the LE/Lys compartment observed upon MOSPD2 or STARD3 loss.

### MOSPD2 requires STARD3 expression to act on lysosomes

To fully establish that LE/Lys homeostasis depends on a functional interplay between MOSPD2 and STARD3, we asked whether MOSPD2 requires STARD3 to restore a normal LE/Lys phenotype. To address this, we repeated the MOSPD2 rescue experiments in cells treated with siRNA targeting or not STARD3. First, as controls, we expressed either the red fluorescent ER marker mScarlet-ER, or mScarlet-tagged MOSPD2 in MOSPD2 KO cells treated with control siRNAs. LE/Lys were then labeled with anti-LAMP1 antibodies and quantified (Fig. 7A, B). Consistent with our previous observations (Fig. 2), MOSPD2 KO cells expressing mScarlet-ER and transfected with control siRNAs exhibited an expanded lysosomal compartment (Fig. 7A, E). As expected, expression of mScarlet-MOSPD2 restored a proper number of LAMP1-positive structures (Fig. 7B, E). Then, we performed the same experiments in cells treated with siRNA targeting STARD3 (Fig. 7C, D); this did not further increase the number of LAMP1-positive vesicles in MOSPD2 KO cells, indicating that the combined absence of STARD3 and MOSPD2 does not worsen the phenotype caused by MOSPD2 deficiency (Fig. 7C, E). Strikingly, when mScarlet-MOSPD2 was re-expressed in MOSPD2 KO cells depleted for STARD3, the number of LE/Lys remained elevated, thus showing that MOSPD2 expression could not correct the phenotype in the absence of STARD3.

**Figure 7:**
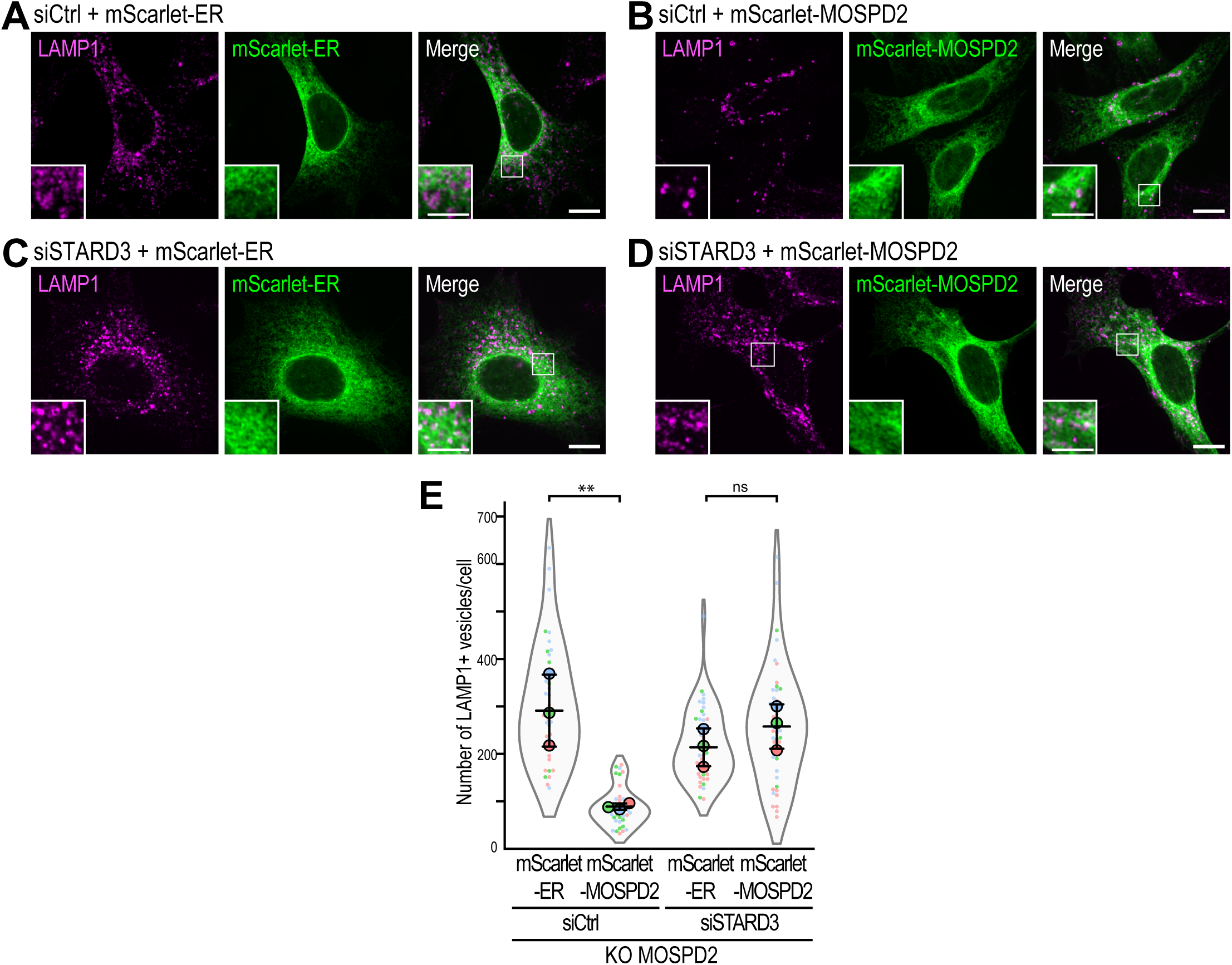
MOSPD2’s activity requires the presence of STARD3. A-D: Confocal images of MOSPD2-deficient HeLa cells transfected with control siRNAs (A, B) or siRNAs targeting STARD3 (C, D) and co-transfected with mScarlet-ER (A, C; green) or mScarlet-MOSPD2 (B, D; green). LE/Lys were stained with anti-LAMP1 antibodies (magenta). E: Quantification of LE/Lys numbers under the conditions shown in panel A-D. Data are displayed as Superplots showing the mean number of LE/Lys per cell (small dots) and per independent experiment (large dots). Independent experiments (n = 3) are color-coded. Means and error bars (SD) are shown as black bars. Data were collected from 39 (siCtrl + mScarlet-ER), 40 (siCtrl + mScarlet-MOSPD2), 46 (siSTARD3 + mScarlet-ER), and 45 (siSTARD3 + mScarlet-MOSPD2) cells. One-way ANOVA with Tukey’s multiple comparisons test (ns, not significant; **, P < 0.01; n = 3 independent experiments).

Overall, these findings indicate that MOSPD2 requires the presence of STARD3 to have a functional impact on LE/Lys, providing functional evidence that MOSPD2 and STARD3 act in the same pathway to regulate these organelles.

### STARD3-mediated cholesterol transport is required for MOSPD2-STARD3 complex function

STARD3 is a cholesterol transporter (Wilhelm et al., 2017; Tsujishita and Hurley, 2000). Given that the STARD3-MOSPD2 complex is required to regulate the sterol content of LE/Lys, in addition to their number and positioning, we investigated whether the cholesterol transport activity of STARD3 was involved in this process. To address this, we performed a structure-function analysis in STARD3 KO cells (STARD3 KO#2) transfected with either the WT or mutated forms of STARD3 with different capacity to bind MOSPD2 (Fig. 8A-H). STARD3 expression was assessed by western blot, as well as its phosphorylation at serine 209 (Fig. 8H). LE/Lys were labeled with Lysotracker, and their cholesterol content assessed using the mEGFP-D4H probe. As previously observed, STARD3 deficiency led to an increased number of LE/Lys strongly labelled with mEGFP-D4H (Fig. 8A-C), accompanied by elevated LAMP2 levels (Fig. 8H). Re-expression of WT STARD3 restored a normal number of LE/Lys, their labeling by the mEGFP-D4H probe (Fig. 8D, I-J), and LAMP2 levels (Fig. 8H). Next, we expressed the STARD3 S_209_A mutant, unable to interact with MSP-domain proteins. This mutant failed to restore either LE/Lys number, cholesterol content, or LAMP2 levels (Fig. 8E, H-J). In contrast, a phosphomimetic mutant of the FFAT motif (STARD3 SD/PA) successfully rescued the phenotype, further supporting that STARD3 needs to bind MOSPD2 to function (Fig. 8F, H-J).

**Figure 8:**
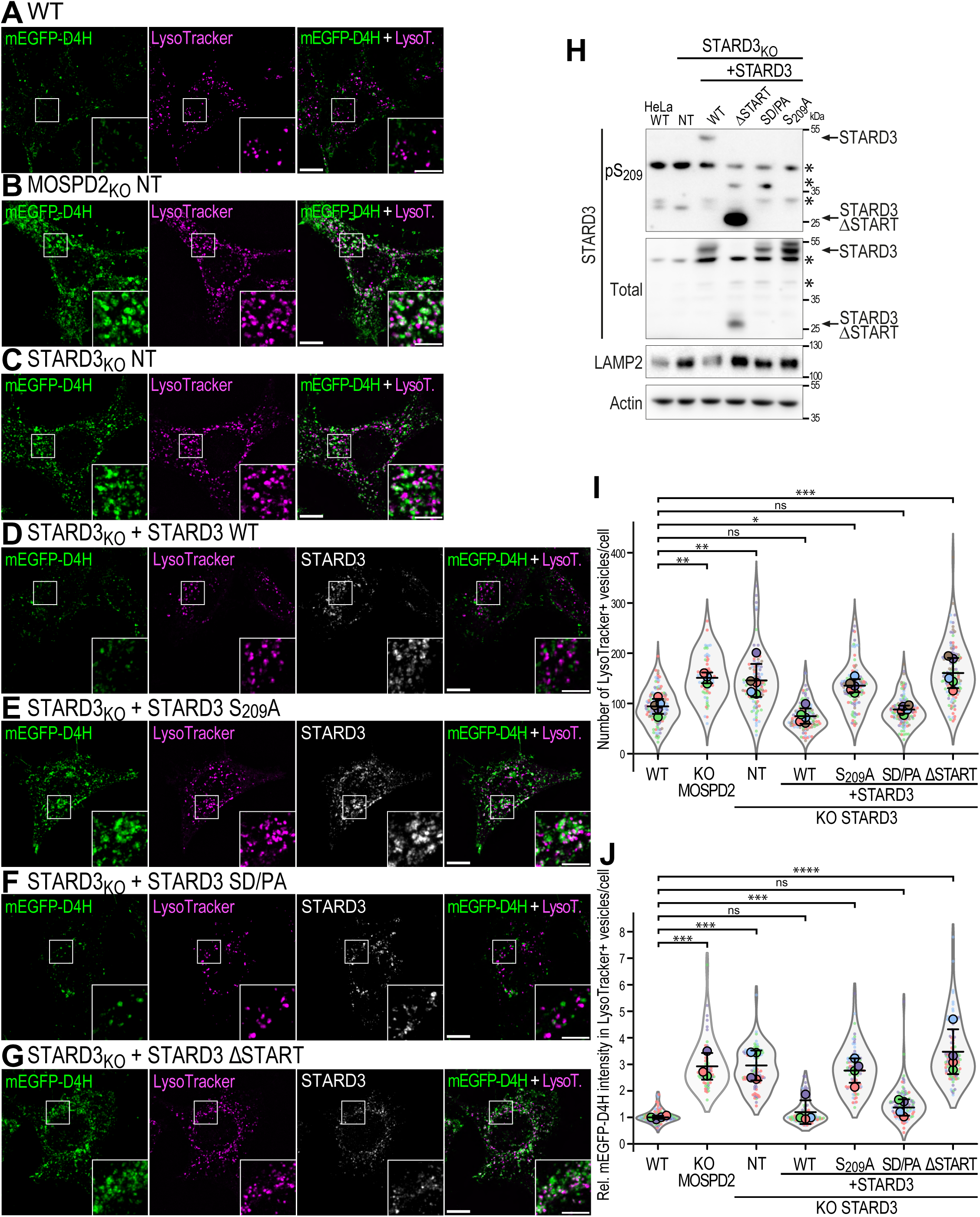
STARD3-mediated cholesterol transport is involved in LE/Lys homeostasis. A-G: Confocal images of WT HeLa cells (A), MOSPD2-deficient HeLa cells (B), and STARD3 KO cells (C-G) either untransfected (A-C) or stably expressing WT STARD3 (D), the STARD3 S_209_A mutant (E), or the STARD3 SD/PA mutant (F), or the STARD3 ΔSTART mutant (G). Cholesterol was labeled with mEGFP-D4H probe (green), LE/Lys with Lysotracker (magenta), and STARD3 with anti-STARD3 antibodies (grey). Higher magnifications images of the area outlined in white are shown on the right. Scale bars: 10 µm; inset scale bars: 4 µm. H: Western Blot analysis of STARD3 protein levels (Total and S_209_ phosphorylation, pS_209_), LAMP2, and Actin in WT HeLa cells and STARD3 KO cells, either untransfected (NT) or transfected with expression vectors encoding WT STARD3, the STARD3 S_209_A mutant, the STARD3 SD/PA mutant, or the STARD3 ΔSTART mutant. I-J: Quantification of LE/Lys number (I) and relative mEGFP-D4H fluorescence intensity in Lysotracker-positive vesicles (J) under the conditions shown in panels A-G. Data are displayed as Superplots showing the mean per cell (small dots) and per independent experiment (large dots). Independent experiments (n = 3-5) are color-coded. Means and error bars (SD) are shown as black bars. Data were collected from 99 (WT), 49 (KO MOSPD2), 85 (KO STARD3 NT), 85 (KO STARD3 + STARD3 WT), 84 (KO STARD3 + STARD3 S209A), 92 (KO STARD3 + STARD3 SD/PA), and 96 (KO STARD3 + STARD3 ΔSTART) cells. One-way ANOVA with Dunnett’s multiple comparisons test (ns, not significant; *, P < 0.05; **, P < 0.01; ***, P < 0.001; ****, P < 0.0001; n = 3-5 independent experiments).

The START domain of STARD3 mediates cholesterol binding and transport between the ER and LE/Lys (Wilhelm et al., 2017; Di Mattia et al., 2020a). We therefore investigated whether it was involved in regulating the sterol content of LE/Lys when STARD3 associates with MOSPD2. To do this, we re-expressed in STARD3 KO cells a mutant lacking the START domain (STARD3 ΔSTART). This mutant was phosphorylated at S_209_ (Fig. 8H) and has previously shown to promote the formation of LE/Lys contacts (Wilhelm et al., 2017). However, this mutant was unable to rescue either the number of LE/Lys, their cholesterol content, or LAMP2 levels (Fig. 8G, H, I).

Together, these findings demonstrate that the cholesterol transport activity of the MOSPD2-STARD3 complex at ER-LE/Lys MCSs is a key regulator or LE/Lys biology.

### STARD3 interacts preferentially with MOSPD2 over VAP-A and VAP-B

VAP-A, VAP-B and MOSPD2 are attached to the ER and project a cytosolic MSP domain to recruit proteins containing an FFAT or a Phospho-FFAT motif. Despite these common properties, our results suggest that they are not functionally interchangeable, as endogenous VAP-A or VAP-B cannot compensate for the loss of MOSPD2 in maintaining LE/Lys homeostasis. We reasoned that this difference might be due to the different capacity of VAP-A, VAP-B and MOSPD2 to recruit specific protein partners, notably STARD3, in cells. In line with this, we previously showed that the MSP domain of MOSPD2 exhibits slight structural differences compared to the MSP domains of VAP-A and VAP-B (Di Mattia et al., 2020a). In addition, we showed that peptides containing the Phospho-FFAT motif sequence of STARD3 bind to the MSP domain of MOSPD2 with 5– to 10-fold higher affinity than to the MSP domains of VAP-A and VAP-B (Di Mattia et al., 2020a). However, these measurements were performed *in vitro* using recombinant MSP domains and short peptides, and may not reflect the behavior of the full-length proteins in the cellular context, where multiple MSP-domain proteins and FFAT-containing partners compete for binding. We therefore sought to determine whether the preferential interaction of STARD3 with MOSPD2 is maintained in the cellular context.

To address this, we first performed immunoprecipitations (Fig. 9A). Flag-tagged STARD3 was co-expressed in HeLa cells with GFP-tagged VAP-A, VAP-B, and MOSPD2. Proteins were then immunoprecipitated using anti-Flag antibodies, and complex formation between STARD3 and VAP-A, VAP-B, or MOSPD2 was analyzed by Western blot (Fig. 9A). As previously described (Di Mattia et al., 2018; Alpy et al., 2013), VAP-A, VAP-B, and MOSPD2 co-immunoprecipitated with STARD3, whereas the MOSPD2 RD/LD mutant, defective in binding FFAT motifs, did not. Importantly, the efficiency of co-immunoprecipitation using STARD3 as a bait varied: MOSPD2 was more efficiently co-precipitated than VAP-A and VAP-B, suggesting that MOSPD2 is the favored partner of STARD3.

**Figure 9:**
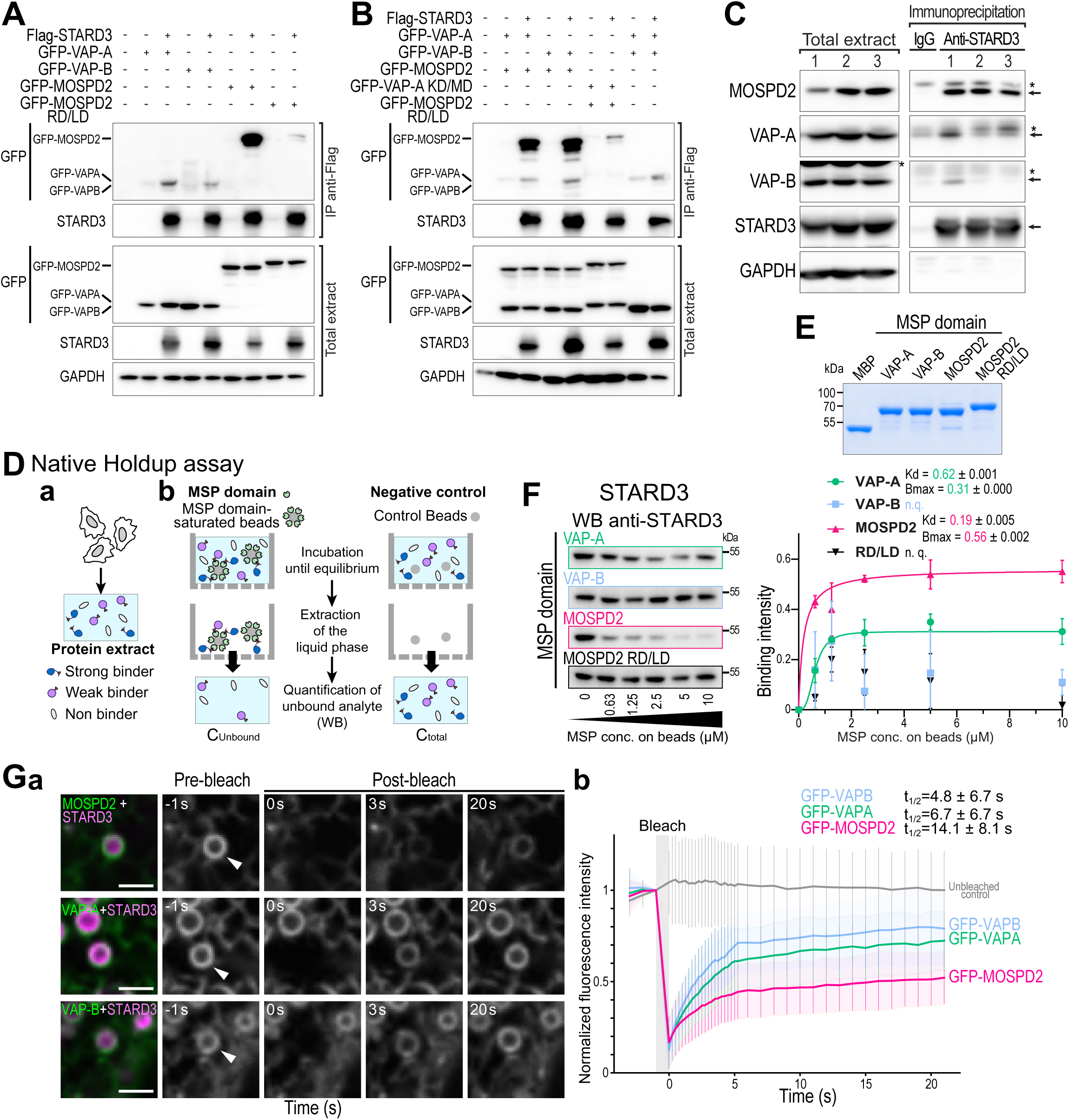
STARD3 binds preferentially to MOSPD2 over VAP-A and VAP-B. A-B: Immunoprecipitation (anti-Flag) experiments between Flag-STARD3 and GFP-VAP-A, GFP-VAP-B, GFP-MOSPD2, GFP-MOSPD2 RD/LD, and GFP-VAP-A KD/MD (B) in HeLa cells. Approximatively 5 µg of total protein extract was analyzed by Western blot using anti-GFP, anti-STARD3, and anti-GAPDH antibodies. Immunoprecipitated proteins were analyzed using anti-GFP and anti-STARD3 antibodies. C: Immunoprecipitation between endogenous STARD3 and VAP-A, VAP-B and MOSPD2 in HCC1954 cells. Immunoprecipitation was performed using control IgG or anti-STARD3 antibodies in triplicate. Total protein extracts and immunoprecipitated proteins were analyzed by Western blot using anti-MOSPD2, anti-VAP-A, anti-VAP-B, anti-STARD3, and anti-GAPDH antibodies. *: aspecific. D: Principle of the native Holdup assay. Total protein extracts (a) are incubated with streptavidin resin saturated with a biotinylated MSP domain or control resin (b). After reaching equilibrium, unbound proteins are filtered out and quantified by Western blot. Binding intensity = 1 – (C_Unbound_ / C_total_). E: Coomassie blue staining of recombinant proteins used for native Holdup experiments: MBP alone or fused to the MSP domains of VAP-A, VAP-B or MOSPD2 (WT and RD/LD mutant), tagged with a 6 His for purification and biotinylated thanks to an AviTag. A total of 25 pmol of each protein was loaded. F: Native Holdup experiments quantifying the interaction between the recombinant MSP domains of VAP-A, VAP-B, MOSPD2, and MOSPD2 RD/LD and endogenous STARD3. Left: western blot analysis of the unbound prey protein (STARD3) in HCC1954 protein extracts after incubation with increasing amounts of the recombinant MSP domains. Right: Binding intensity between the MSP domains and the prey protein (STARD3). Binding curves were fitted using a Hill equation (mean ± SEM from 2 technical replicates), and apparent affinities (*K*_app_) and maximal binding intensities (*B*_max_) were calculated (± SD). G: FRAP experiment in HeLa cells co-expressing mCherry-STARD3 and either GFP-VAP-A, GFP-VAP-B, or GFP-MOSPD2. a: images showing STARD3-positive LE/Lys in close apposition to ER-localized GFP-MOSPD2 (top), GFP-VAP-A (middle), or GFP-VAP-B (bottom). Left: colocalization of mCherry-STARD3 (magenta) and GFP (green) pre-bleach. GFP signal (gray) displayed sequentially from left to right: pre-bleach, immediately post-bleach, 3 seconds post-bleach, and 20 s post-bleach. b: Quantification of relative GFP-signal intensity in the bleached ROI during the 2 seconds before bleaching, and the 21 seconds following bleaching in cells expressing GFP-VAP-A (blue curve), GFP-VAP-B (green curve), and GFP-MOSPD2 (red curve). The gray curve represents the GFP signal in the absence of bleaching. Mean values and standard deviations (black bars) are shown. The calculated half-times of recovery (mean t½ ± SD) are indicated.

Then, to further analyze the ability of STARD3 to selectively interact with MOSPD2, we set up immunoprecipitation assays based on competition (Fig. 9B). To this end, we established stable cell lines expressing different combinations of GFP-tagged proteins: GFP-MOSPD2 with GFP-VAP-A, GFP-MOSPD2 with GFP-VAP-B, and GFP-VAP-A with GFP-VAP-B. As a negative control, we also generated a cell line expressing two mutants unable to bind FFAT motifs, GFP-VAP-A KD/MD and GFP-MOSPD2 RD/LD. Flag-tagged STARD3 was expressed in the different cell lines and then immunoprecipitated. The amounts of GFP-VAP-A, –VAP-B and –MOSPD2 co-immunoprecipitated with STARD3 were assessed by Western blot. Although the levels of GFP-tagged proteins were similar in the cell extracts, more GFP-MOSPD2 was co-immunoprecipitated with Flag-STARD3 than GFP-VAP-A and GFP-VAP-B. Of note, GFP-VAP-A and GFP-VAP-B were only weakly co-immunoprecipitated with STARD3, even in cells expressing both proteins without GFP-MOSPD2 overexpression. These results confirm that STARD3 binds more efficiently to MOSPD2 than to VAP-A or VAP-B.

Finally, to validate our findings in a more physiological setting, we investigated the interaction between endogenous STARD3 and its binding partners. To this end, we used HCC1954 breast cancer cells, which endogenously express high levels of STARD3 (Fig. 9C). STARD3 was immunoprecipitated using an anti-STARD3 antibody, and VAP-A, VAP-B, and MOSPD2 were detected by Western blot. While MOSPD2 was efficiently immunoprecipitated, only small amounts of VAP-A and VAP-B were recovered in the immunoprecipitated fractions. Collectively, our data support the notion that STARD3 preferential bind to MOSPD2 in a cell context.

### Quantification of the specific MOSPD2-STARD3 interaction in cells

To further confirm STARD3’s preference for MOSPD2 in a cellular context, we quantified the binding affinities of MOSPD2, VAP-A, and VAP-B for endogenous STARD3 using native Holdup assays. This method quantifies protein-protein interaction affinities by measuring the extent to which a prey protein is depleted from a cell lysate after reaching binding equilibrium with an immobilized bait under native conditions (Zambo et al., 2022; Delalande et al., 2025; Gogl et al., 2022) (Fig. 9D).

To perform these assays, we used as baits the recombinant MSP domains of VAP-A, VAP-B or MOSPD2 (WT and RD/LD mutant) fused to the Maltose Binding Protein (MBP) and tagged with a 6 His for purification. These recombinant proteins were produced in bacteria with an AviTag that allowed *in vivo* biotinylation, purified (Fig. 9E), and immobilized on streptavidin beads. As cell lysate, we used protein extracts from HCC1954 cells which express a number of FFAT-containing proteins including OSBP, ORP1L and STARD3. To assess binding affinities, we performed titrations using increasing concentrations of immobilized MSP domains (Fig. 9F). Protein extracts were incubated with the immobilized MSP domains, and after binding equilibrium was reached, the amount of unbound STARD3 was quantified by western blot. A gradual depletion of STARD3 in the unbound fractions was evident for the MSP domain of MOSPD2, markedly reduced for the MSP domain of VAP-A, and not significant for the MSP domain of VAP-B (Fig. 9F). Quantification of the apparent dissociation constant (*K*_d_) and of the maximal binding capacity (*B*_max_) showed that MOSPD2 binds STARD3 with higher affinity than VAP-A (MOSPD2: *K*_d_ = 0.19 µM, *B*_max_ = 0.56; VAP-A: *K*_d_ = 0.62 µM, *B*_max_ = 0.31). Binding to VAP-B was too weak to reliably determine *K*_d_ or *B*_max_. These results indicate that MOSPD2 interacts more with STARD3 than VAP-A or VAP-B do, suggesting that it is the predominant binding partner of STARD3.

Next, to examine whether other FFAT-containing proteins show similar binding preferences, we assessed the interactions of MOSPD2, VAP-A, and VAP-B MSP domains with endogenous ORP1L and OSBP (Fig. S4). Interestingly, MOSPD2 was strongly bound to ORP1L, but did not interact with OSBP. OSBP behaved differently than STARD3 and ORP1L, as it was predominantly bound to VAP-A and VAP-B. Together, these results demonstrate that MOSPD2, VAP-A, and VAP-B have distinct binding affinities for different FFAT-containing partners. This selectivity indicates that FFAT-mediated interactions are not interchangeable, but instead define specific protein-protein interaction networks.

To further investigate the differential binding of STARD3 to the various VAP proteins in living cells, we performed Fluorescence Recovery After Photobleaching (FRAP) experiments. We imaged cells co-expressing mCherry-tagged STARD3 with GFP-tagged VAP-A, VAP-B, or MOSPD2 (Fig. 9G). As expected, VAP-A, VAP-B and MOSPD2 accumulated around STARD3-positive LE/Lys (Fig. 9Ga). GFP fluorescence around individual LE/Lys was bleached and the fluorescence recovery was measured (Fig. 9Gb). The fluorescence recovery of GFP-VAP-A and GFP-VAP-B was similar (t_1/2_ of ∼7 and ∼5 s, respectively), indicating that the exchange of these two proteins between areas of the ER in contact with STARD3-positive vesicles and the rest of the ER occurs at similar rate. In contrast, GFP-MOSPD2 showed a slower fluorescence recovery (t_1/2_ of ∼14 s), suggesting that the protein is more stably associated with STARD3 at ER-LE/Lys contacts than VAP-A and VAP-B. Thus, in live cells, MOSPD2 associates more stably with STARD3 than VAP-A and VAP-B do.

Together, these data show that, in a cell context, STARD3 preferentially forms complexes with MOSPD2 over VAP-A and VAP-B, and more broadly, that VAP proteins exhibit distinct binding preferences for different FFAT-containing partners.

### STARD3 overexpression overrides the loss of MOSPD2

According to the Protein Abundance Database (PaxDb; Dataset Human PeptideAtlas; whole organism, integrated), STARD3 is a low abundance protein (0.32 ppm) compared to VAP-A (154 ppm), VAP-B (53 ppm) and MOSPD2 (1.43 ppm) (Huang et al., 2023). We reasoned that increasing the concentration of STARD3 might promote the formation of alternate protein complexes with VAP-A and VAP-B, even though STARD3 has a lower affinity for these two proteins, and that this could potentially compensate for the absence of MOSPD2. To test this hypothesis, we transfected MOSPD2-deficient cells with a STARD3-expressing construct and analyzed the LE/Lys phenotype using LAMP1 labelling (Fig. 10). Compared with untransfected MOSPD2-deficient cells, cells with STARD3 overexpression showed a reduced number of LAMP1-positive structures; reaching levels comparable to those observed in WT cells (Fig. 10A-C and G).

**Figure 10:**
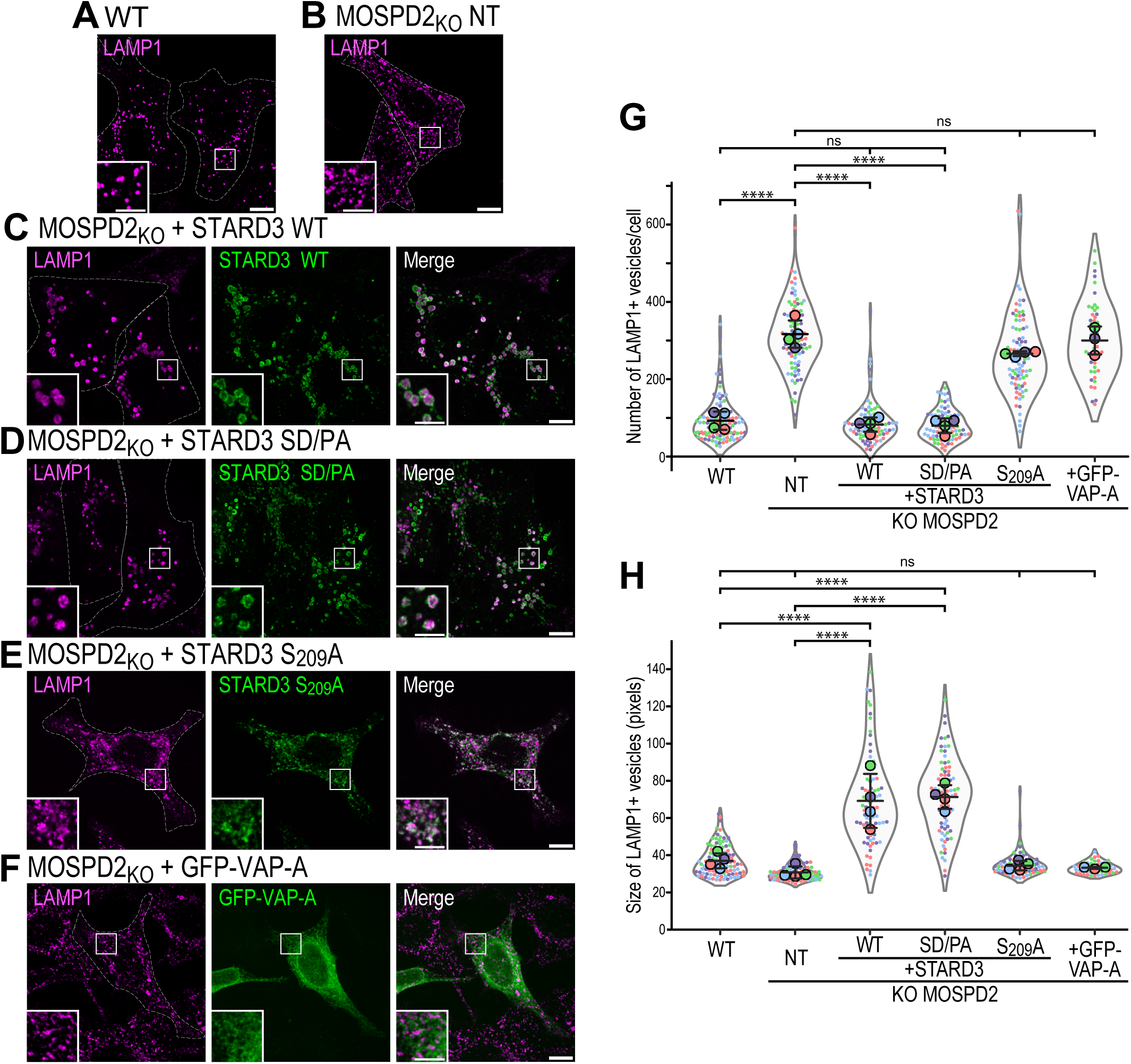
The overexpression of STARD3 rescues the LE/Lys phenotype in MOSPD2-deficient cells. A-F: Confocal images of WT HeLa cells (A) and MOSPD2-deficient HeLa cells (B-F), either untransfected (A-B) or transfected with expression vectors encoding WT STARD3 (C), the STARD3 SD/PA mutant (D), the STARD3 S209A mutant (E), or GFP-VAP-A (F). LE/Lys were labeled with anti-LAMP1 antibodies (magenta), and STARD3 was detected using anti-STARD3 antibodies. G-H: Quantification of LE/Lys numbers (G) and sizes (H) under the conditions shown in panels A-F. Data are displayed as Superplots showing the mean per cell (small dots) and per independent experiment (large dots). Independent experiments (n = 3-4) are color-coded. Means and error bars (SD) are shown as black bars. Data were collected from 108 (WT), 95 (KO MOSPD2 NT), 86 (KO MOSPD2 + STARD3 WT), 84 (KO MOSPD2 + STARD3 SD/PA), 86 (KO MOSPD2 + STARD3 S209A), and 48 (KO MOSPD2 + VAP-A) cells. One-way ANOVA with Tukey’s multiple comparisons test (ns, not significant; ****, P < 0.0001; n = 3-4 independent experiments).

The interaction of STARD3 and the MSP domain requires the phosphorylation of its Phospho-FFAT on S209 residue. The phosphomimetic mutation S_209_D/P_210_A (thereafter named SD/PA) in STARD3 promotes its stable association with the MSP domain, whereas the S_209_A mutation, which prevents its phosphorylation, impedes its binding with MODP2, VAP-A, and VAP-B (Di Mattia et al., 2020a). We used these mutants to test whether the ability of STARD3 to overcome the loss of MOSPD2 needs a functional FFAT motif and interaction with VAP-A or VAP-B. Of interest, the phosphomimetic STARD3 SD/PA mutant restored a normal LE/Lys phenotype in MOSPD2-deficient cells, similarly to WT STARD3 (Fig. 10D, G). In contrast, the STARD3 S_209_A mutant did not rescue the phenotype, indicating that a functional FFAT motif, and thus binding to MSP domain proteins, is required for overexpressed STARD3 to compensate for the loss of MOSPD2, most likely through interactions with other members of the VAP family, i.e. VAP-A and VAP-B. Of note, the overexpression of WT or SD/PA mutant STARD3 resulted in an increase in LE/Lys size (Fig. 10H), as observed in previous studies (Alpy et al., 2005; Holtta-Vuori et al., 2005). Finally, we overexpressed VAP-A in MOSPD2-deficient cells and we observed that it did not compensate for the absence of MOSPD2, further suggesting that the amount of STARD3 is the limiting factor (Fig. 10F, G).

Overall, these data show that STARD3 interacts with its VAP partners with different affinities. In WT cells, where STARD3 has a low abundance, the STARD3-MOSPD2 complex is preferentially formed and functions to regulate LE/Lys. Increasing STARD3 levels allows it to bypass the requirement for MOSPD2 by engaging alternative VAP partners, thereby restoring proper LE/Lys homeostasis.

## Discussion

In this study, we identify MOSPD2, an ER-anchored protein, as a new key regulator of LE/Lys homeostasis. We show that this function relies on the selective association of MOSPD2 with the LE/Lys cholesterol transfer protein STARD3. Inactivation of either MOSPD2 or STARD3 results in a similar phenotype characterized by an expansion of the LE/Lys compartment, cholesterol accumulation at LE/Lys, and impaired fusion capacity. These findings indicate that MOSPD2 and STARD3 operate in a common functional pathway at ER-LE/Lys contact sites. Furthermore, we establish that although MOSPD2 has many protein partners, it forms a functional and privileged complex with STARD3; this interaction is direct and requires a binding-competent MSP domain in MOSPD2 and a functional FFAT motif in STARD3. Reciprocally, we found that STARD3 preferentially associates with MOSPD2 rather than with its homologues VAP-A and VAP-B. Moreover, we show that VAP-A and VAP-B cannot compensate for the loss of MOSPD2, and other LE/Lys-associated cholesterol transport proteins, such as ORP1L, cannot compensate for the loss of STARD3. Collectively, these findings indicate that MOSPD2 and STARD3 constitute a unique functional complex at ER-LE/Lys contact sites.

The ER possesses a small group of MSP-domain receptors (VAP-A, VAP-B and MOSPD2) that mediate the formation of ER-organelle contacts by binding FFAT– or Phospho-FFAT-containing partners on opposing organelle membranes or the plasma membrane. Proteomics databases list hundreds of potential binding partners for these proteins, many of which are shared, implying that FFAT-containing partners likely compete for binding. This observation raises the central question of how MSP-domain proteins select in a hierarchical manner their partner proteins to control key cellular processes. Several parameters are expected to shape the “interaction landscape” of MSP-domain proteins: First, intrinsic protein features, such as the unique identity and surface properties of each MSP domain, the primary sequence of FFAT motifs, and their post-translational modifications, notably phosphorylation, determine the affinity between MSP-domain proteins and their FFAT-containing partners. Second, the cellular context in which these interactions occur, including the relative abundance of MSP-domain and FFAT-containing proteins, their local enrichment, as well as potential conformational changes, controls which interactions can actually form *in vivo*.

Our data uncover some of these parameters. We previously showed that synthetic peptides containing the Phospho-FFAT motif sequence of STARD3 bind *in vitro* to the MSP domain of MOSPD2 with 5– to 10-fold higher affinity than to the MSP domains of VAP-A and VAP-B (Di Mattia et al., 2020a). Here, we extend these measurements to the cellular context using the Holdup assay, and show that the MSP domain of MOSPD2 binds STARD3 with a higher affinity than the MSP domains of VAP-A and VAP-B. This implies that this interaction is determined both by the specific features of the MOSPD2 MSP domain and by the sequence and post-translational modifications of the STARD3 FFAT motif, and highlights binding affinity as a key principle in organizing MSP-mediated interaction networks. While the MSP domains of VAP-A and VAP-B are closely related (82% identity), the MSP domain of MOSPD2 is more divergent (28% identity with VAP-A and 30% with VAP-B). Although the overall fold of the MSP domain is conserved, discrete variations at the binding interfaces modulate interactions with a given FFAT-containing protein (Di Mattia et al., 2020a; Kaiser et al., 2005; Shi et al., 2010).

Second, we provide data showing that variations in FFAT motifs presumably fine-tune binding affinities in cells. It has been predicted that numerous FFAT motifs corresponds to slight or more important variations of the canonical FFAT consensus sequence (EFFDAxE) and that this presumably modulates their affinity for MSP domains (Levine, 2025). Consistent with this view, we show here using a semi-quantitative *in vitro* binding assay that FFAT-containing proteins have specific binding profiles: for example, OSBP interacts with VAP-A and VAP-B but poorly with MOSPD2, whereas ORP1L binds more strongly to MOSPD2 than to VAP-A or VAP-B. These observations suggest that FFAT-containing proteins differ in their binding to MSP-domain tethers, exhibiting selective interaction patterns that favor one or more specific partners. As a consequence, competition among FFAT-containing proteins implies that even small differences in binding affinity may have major functional consequences. Changes in affinity could determine whether a given FFAT-containing protein can engage an MSP-domain partner or is instead outcompeted by others. Importantly, FFAT motifs consist of a core sequence flanked by an acidic tract, and the number of negative charges within this region, including those introduced by phosphorylation, correlates with binding affinity for MSP domains (Levine, 2025; Di Mattia et al., 2020a). This highlights the crucial regulatory role of phosphorylation to control the activity of FFAT-containing proteins, and more broadly the dynamics and function of MCSs. Consistent with this idea, phosphorylation of the acidic tract of the FFAT motif in the ceramide transporter CERT (also known as STARD11) enhances its interaction with VAP and stimulates ceramide transport at ER-Golgi contact site (Kumagai et al., 2014).

Other more contextual parameters are also likely major determinants that define the landscape of interactions at MCSs. According to the Protein Abundance Database (Huang et al., 2023) and our previous study in Hela cells (Di Mattia et al., 2018), VAP-A is substantially more abundant than VAP-B, which itself is more abundant than MOSPD2. Thus, according to the law of mass action, for interactions of comparable affinity, VAP-A is expected to form more complexes with FFAT-containing proteins. This is consistent with physiological data in mice, where VAP-A is essential for survival, while VAP-B– and MOSPD2-deficient mice remain viable (McCune et al., 2017; Kabashi et al., 2013; Yacov et al., 2020). In this context, our data show that the weak interactions between STARD3 and VAP-A or VAP-B, which are insufficient to elicit a biological effect under normal conditions, can be partially compensated by increasing STARD3 expression. Importantly, these findings establish that MSP-domain proteins are not necessarily functionally interchangeable for all FFAT-containing partners. In the case of STARD3, the combination of high binding affinity for MOSPD2 and lower affinity for VAP proteins results in a non-redundant requirement for MOSPD2 despite its lower concentration, whereas for other FFAT-containing proteins with similar affinities for multiple MSP-domain proteins and/or higher expression levels, MSP-domain proteins may be interchangeable.

Finally, the local enrichment of MSP-domain tethers within the ER is also likely to influence the dynamics of their interactions and, consequently, their functional outcomes. While the localization of overexpressed MSP-domain proteins is often shaped by the formation of MCSs (Di Mattia et al., 2018; Obara et al., 2024; Alpy et al., 2013), the spatial distribution of endogenous MSP-domain proteins remains poorly understood. It is still unclear which fraction of these proteins is uniformly distributed throughout the ER and which fraction is instead enriched in discrete ER subdomains. In addition, conformational regulation of FFAT-containing proteins provides another layer of control over MSP-domain interactions. For several of them, such as ORP1L, intramolecular interactions can hide the FFAT motif, and release of this autoinhibited state, triggered by changes in lipid binding, exposes the motif and promotes binding to MSP-domain proteins (Rocha et al., 2009). This illustrates how conformational switches can dynamically regulate when and where MSP-mediated contacts are formed.

MSP-domain proteins found in mammals are paralogs. Proteomic studies indicate that they share highly similar interactomes, both in the number and identity of their binding partners, suggesting that they might be functionally redundant. However, standard proteomic approaches do not provide information on the relative binding affinities or competitive hierarchies between partners, and therefore cannot predict which interactions will actually dominate in a cellular context. Yet, some functional studies support this redundancy notion; for example, VAP-A and VAP-B have a redundant function in autophagy (Mao et al., 2019) and on bidirectional traffic between endosomes and the Golgi (Dong et al., 2016). However, other functional studies show that it is not always the case: for example, we reported a unique function for MOSPD2 in lipid droplet biology (Zouiouich et al., 2022), and VAP-A was specifically implicated in mitochondria homeostasis (Subra et al., 2023a). In this study, we provide another example of a specific function of MOSPD2 on LE/Lys biology. Moreover, we show that this specificity is not held by MOSPD2 alone but is shared with one of its partners STARD3 supporting the idea that the complex formed by MOSPD2 and STARD3 at ER-LE/Lys MCS is implicated in LE/Lys biology. Given that STARD3 is a sterol transport protein and that LE/Lys MCS are active zone of cholesterol exchange (Ikonen and Olkkonen, 2023), we investigated the impact of this specific interaction of cholesterol distribution.

Our results show that disrupting the MOSPD2-STARD3 complex leads to the accumulation of free cholesterol at LE/Lys, indicating that this complex contributes to sterol distribution and LE/Lys homeostasis. Consistent with this, an unbiased screen using the GFP-D4 probe identified MOSPD2 and STARD3 among factors whose silencing increases LE/Lys sterol content (Lu et al., 2021). Puzzlingly, several studies have reported that STARD3 overexpression can also result in cholesterol accumulation in LE/Lys (Wilhelm et al., 2017; Holtta-Vuori et al., 2005; Liapis et al., 2012), indicating that STARD3 function is not simply proportional to its abundance: STARD3 when insufficient or in excess similarly impacts cholesterol homeostasis. How to explain this apparent paradox? At endogenous levels, STARD3 likely participates in regulated sterol exchange at ER-LE/Lys contact sites, and its loss may impair cholesterol flux out of LE/Lys, leading to sterol accumulation. In contrast, STARD3 overexpression, by promoting excessive ER-LE/Lys contacts, may induce non-physiological sterol accumulation and, subsequently, sterol-dependent membrane remodeling. In support of this idea, STARD3 overexpression induces not only cholesterol accumulation but also the buildup of intraluminal membranes within LE/Lys (Wilhelm et al., 2017), a phenotype that we did not observe upon MOSPD2 or STARD3 depletion. This suggests that STARD3 overexpression mediate parallel sterol fluxes, in addition of the endogenous function of the MOSPD2-STARD3 complex. Together, these observations support a model in which STARD3 activity must be tightly regulated, and in which proper recruitment of STARD3 by MOSPD2 at defined ER-LE/Lys contact sites is required for balanced cholesterol homeostasis.

Cholesterol levels at LE/Lys membrane are known to regulate the positioning of these organelles (Cabukusta and Neefjes, 2018). Increased cholesterol has been reported to drive perinuclear clustering of LE/Lys through an ORP1L-dependent mechanism (Rocha et al., 2009). In contrast, we observed that, in STARD3– and MOSPD2-deficient cells, cholesterol accumulation correlates with a more peripheral distribution of LE/Lys. This suggest that STARD3 operates independently of ORP1L consistent with their localization to distinct LE/Lys sub-populations (Kant et al., 2013). In addition to positioning defects, we also observed that MOSPD2 KO cells display defects in LE/Lys fusion capacity. Similarly, and consistent with earlier studies (Holtta-Vuori et al., 2005), STARD3 KO cells also display fusion impairments. The elevated cholesterol levels in LE/Lys of MOSPD2– or STARD3-deficient cells points to a potential link between sterol accumulation in LE/Lys and fusion defects. Indeed, cholesterol is known to modulate membrane biophysical properties and influence membrane fusion (Yang et al., 2016), providing a plausible mechanistic explanation for the fusion defects observed upon loss of MOSPD2 or STARD3.

In summary, our results indicate that MSP-domain proteins do not simply act as generic membrane tethers. By selectively interacting with specific FFAT-containing partners, they help shape the molecular composition and function of individual MCSs. The MOSPD2-STARD3 complex at ER-LE/Lys contacts illustrates this principle, revealing a previously unrecognized role for these contacts in LE/Lys homeostasis and cholesterol distribution. Such selectivity provides a way for cells to control organelle communication and regulate their functions.

## Materials and Methods

### Cloning and constructs

The pQCXIP MOSPD2 (WT; RD/LD; ΔCRAL-TRIO; ΔMSP), pQCXIP GFP-MOSPD2 (WT: Addgene #186467; RD/LD #186468), pQCXIP mScarlet-MOSPD2 (WT: Addgene #186472; RD/LD Addgene #186476; W201E: Addgene plasmid #186477; ΔMSP; ΔCRAL-TRIO mutants), GFP-VAP-A (WT: Addgene #104447; KD/MD Addgene #104449), GFP-VAP-B (Addgene plasmid #104448), EGFP-ER[TM(SAC1)] (Addgene plasmid #186475), pQCXIP mScarlet-ER [TM(SAC1)] (Addgene plasmid # 186572), Flag-STARD3, GFP-VAP-A (WT and KD/MD mutant), and GFP-VAP-B, expression vectors were previously described (Alpy et al., 2005, 2013; Di Mattia et al., 2018; Zouiouich et al., 2022; Eichler et al., 2026). The pLKO.1 vectors encoding shRNAs targeting MOSPD2 were also reported earlier (Di Mattia et al., 2018). EGFP-D4 and mEGFP-D4H expression constructs have been described previously (Eichler et al., 2025). The mCherry-STARD3 expression vector was described in (Alpy et al., 2013).

pQCXIP STARD3, pQCXIP STARD3 S_209_D/P_210_A, and pQCXIP STARD3 S_209_A retroviral expression vectors were described previously (Alpy et al., 2013; Di Mattia et al., 2020a). pLenti PGK Puro^R^ encoding STARD3 (WT, SD/PA, ΔSTART, S_209_A) were previously described (Eichler et al., 2026).

The pQCXIP vector was modified by replacing the puromycin resistance gene with a blasticidin resistance gene. To this end, two MfeI restriction sites were first introduced by successive site-directed mutagenesis (QuickChange, Agilent) using the primers 5’-CCA CAA GGA GAC GAC CAA TTG TGA CCG AGT ACA AGC C-3’ and 5’-CCC GCA AGC CCG GTG CCA ATT GCC CGC CCC ACG ACC CGC-3’. The blasticidin resistance gene was then amplified from pLenti PGK Blast^R^ (Eichler et al., 2026) by PCR using the primers 5’-GAG AGA ATT CGG AAC TAA ACC ATG GCC AAG C-3’ and 5’-GAG AGA ATT CTT AGC CCT CCC ACA CAT AAC C-3’. The resulting PCR fragment was digested with EcoRI and inserted into the modified pQCXIP vector linearized with MfeI to generate the pQCXIP Blast^R^ vector. The coding sequence of VAP-A (WT and KD/MD mutant) and VAP-B fused to EGFP were cloned by SLiCE into the pQCXIP Blast^R^ vector linearized with NotI and BamHI. Inserts were amplified by PCR using the following primers: 5’-AAT TGA TCC GCG GCC ACC ATG GTG AGC AAG GGC GAG-3’ and 5’-GCG GAA TTC CGG ATC GCT ACA AGA TGA ATT TCC CTA GAA AGA ATC CAA TG-3’ for VAP-A, and 5’-GCG GAA TTC CGG ATC GCT ACA AGG CAA TCT TCC CAA TAA TTA CAC-3’ for VAP-B.

pLenti PGK mEGFP-D4H was generated by SLiCE cloning of a PCR-amplified mEGFP-D4H fragment (Eichler et al., 2025) using primers: 5‘-AGGGG GATCC ACCGG ACCAG GATGG TGAGC AAGGG CGAGG AGCTG TTCAC CGGGG TGGTG CCC-3’ and 5‘-TCAAC CACTT TGTAC ATTAA TTGTA AGTAA TACTA GATCC AGGGT ATAAA GTTGT TC-3’ into AgeI– and BsrGI-linearized pLenti PGK Puro^R^.

Bacterial expression constructs consisting of a fusion between the Avitag, a His6 tag, MBP (Maltose Binding Protein), a TEV cleavage site, and the MSP domain of VAP-A, VAP-B, or MOSPD2 (WT and RD/LD mutant) were generated. To this aim, PCR fragments encoding the MSP domain of VAP-A, VAP-B, and MOSPD2 were obtained using the following primers: VAP-A: 5’-TTTCA GGGCG CCATG GCGAA GCACG AGCAG ATCCT GGTCC TCG-3’ and 5’-GACCG TTATA GTTAG TCACA ATTTA TCATT TTCAT TGGGC ATTTC AAATA CGC-3’; VAP-B: 5’-TTTCA GGGCG CCATG GCGAA GGTGG AGCAG GTCCT GAGCC-3’ and 5’-GACCG TTATA GTTAG TCATG GTTTA TCATT CTCTG CTGGC AATTC AAACA C-3’; MOSPD2: 5’-TTTCA GGGCG CCATG ATTGC TTTCA AGAAA CCATT GAGTG TATTT AAAGG CCC-3’ and 5’-GACCG TTATA GTTAG TCAAA CAGTA TGGCA TCTTA ACCTA TGTTC CATC-3’ and pQCXIP VAP-A, pQCXIP VAP-B, pQCXIP MOSPD2 and pQCXIP MOSPD2 RD/LD as templates, respectively. PCR fragments were cloned by SLiCE into the NcoI and KpnI linearized pETM41a vector (EMBL, Heidelberg, Germany).

All constructs were verified by DNA sequencing (Eurofins).

### Cell culture, transfection and infection

HeLa cells [American Type Culture Collection (ATCC) CCL-2, RRID:CVCL_0030] were maintained in DMEM (1 g/L glucose) with 5% fetal calf serum (FCS) and 40 µg/mL gentamycin. 293T cells (ATCC CRL-3216) were maintained in DMEM (1 g/L glucose) with 10% FCS, 100 UI/mL penicillin and 100 µg/mL streptomycin. MRC5 cells (ATCC CCL-171) were maintained in DMEM (1 g/L glucose) with 10% FCS and 40 µg/mL gentamycin. HCC1954 cells (ATCC CRL-2338, RRID:CVCL1259) were maintained in RPMI 1640 without HEPES with 10% FCS and 40 µg/mL gentamycin.

Cells were transfected with X-tremeGENE 9 DNA Transfection Reagent (Roche) following the manufacturer’s instructions.

To generate retroviral particles, pQCXIP vectors were co-transfected with pCL-Ampho vector (Imgenex) into the 293T retroviral packaging cell line. The resulting viral particles, supplemented with 10 µg/mL polybrene and 20 mM HEPES, were then incubated with the recipient cells. Stable cells were selected with puromycin (0.5 µg/mL), blasticidin (4 µg/mL), or both, depending on the resistance markers of the vectors used. Retroviral infections were used to generate HeLa/Ctrl, HeLa/GFP-MOSPD2, HeLa/GFP-MOSPD2 RD/LD, HeLa/GFP-MOSPD2 ΔCC, HeLa/GFP-MOSPD2 RD/LD ΔCC. HeLa/GFP-MOSPD2 ΔMSP, HeLa/GFP-MOSPD2 W201E, HeLa/GFP-MOSPD2 ΔCRAL-TRIO, and HeLa/mScarlet-ER cell lines. The HeLa/Ctrl cell line was obtained using the empty pQCXIP plasmid. To generate double stable cell lines, HeLa/GFP-MOSPD2 cells (puromycin-resistant) were infected with pQCXIP Blast^R^ GFP-VAP-A or pQCXIP Blast^R^ GFP-VAP-B, HeLa/GFP-MOSPD2 RD/LD cells (puromycin resistant) were infected with pQCXIP Blast^R^ GFP-VAP-A KD/MD, and HeLa/GFP-VAP-B cells (puromycin resistant) were infected with pQCXIP Blast^R^ GFP-VAP-A.

To generate lentiviral particles, pLenti PGK Puro^R^ vectors were co-transfected with three packaging plasmids (pLP1, pLP2, and pLP/VSVG from Invitrogen) into 293T cells. Lentiviral transduction was used in STARD3 KO cells (clone #2) to generate HeLa KO STARD3/STARD3 WT, HeLa KO STARD3/STARD3 ΔSTART, HeLa KO STARD3/STARD3 S209A, and HeLa KO STARD3/STARD3 SD/PA cell lines. Lentiviral transduction of HeLa cells was used to generate the HeLa/mEGFP-D4H reporter cell line.

For siRNA experiments, 3.0 x 10^5^ cells or 1.0 x 10^6^ cells were seeded in 6-well plates or 100-mm dishes, respectively. siRNA transfections were performed using Lipofectamine RNAiMAX (Invitrogen) according to the manufacturer’s instructions. The culture medium was changed the day after transfection, and cells were analyzed 3 days post-transfection. For immunofluorescence (IF) experiments following siRNA transfection, cells were replated into 24-well plates containing glass coverslips the day after transfection, and IF was performed 3 days post-transfection. Control siRNA (D-001810-10) and siRNA targeting VAP-A (L-021382-00-0010), VAPB (L-017795-00-0010), MOSPD2 (L-017039-01-0010), STARD3 (L-017665-00-0010), OSBP (L-009747-00-0005), OSBPL1A (L-008350-00-0005) were SMARTpool ON-TARGETplus obtained from Horizon Discovery.

### CRISPR/Cas9-mediated genome editing

To generate STARD3 KO clones, HeLa cells were plated in 100 mm dishes and transfected with an expression vector (pX330 Cas9 HF-T2A-GFP) encoding the Cas9 HF nuclease and 2 gRNAs using X-tremeGENE 9 DNA Transfection Reagent (Roche). The guide RNAs target STARD3 exon 5 and 6 using the sequences: 5’-GTC TGC AGC AGG ATA AAC TG-3’; 5’-TCC TTC TAT CAG GCT CCC TG-3’). 48 hours after transfection, GFP-positive clones were sorted and isolated in 96-well plates using fluorescence-activated cell sorting (FACS, Fusion). Clones were then screened by PCR (5’-GGA GAG GGA ATC CTG TCC TTT GGT ATC-3’ and 5’-ATC TGA TTC ATT GTC AGA CCC TGC AG-3’) and the PCR fragments sequenced by the Sanger method (Eurofins). Positive clones were further analyzed by Western Blot (rabbit anti-STARD3, 1/1000, 605, IGBMC).

MOSPD2-deficient HeLa cells and HeLa cells in which mClover3 was inserted at the endogenous MOSPD2 locus were previously described (Zouiouich et al., 2022).

### Cell lysis

#### For GFP-trap assays

Cells were washed twice with 1x TBS and scraped in ice-cold IP Buffer (50 mM Tris pH 7.5, 50 mM NaCl, 1% Triton X-100, 1 mM EDTA) supplemented with protease (cOmplete, Roche) and phosphatase (PhosSTOP, Roche) inhibitor tablets. Lysates were incubated on ice for 20 min and centrifuged at 9,500 x *g* for 10 min at 4 °C to remove debris. Supernatant were collected, and protein concentration was determined using the BCA protein assay (ThermoFisher Scientific).

#### For immunoprecipitation assays

The procedure was identical to that described above, except that the IP buffer was replaced with ice-cold M-PER buffer (78501, ThermoFisher Scientific) supplemented with cOmplete and PhosSTOP.

#### For native Holdup assays

HCC1954 cells (3.0 x 10^6^) were seeded in 150-mm dishes. At confluence, cells were processed as described above using ice-cold Holdup buffer (50 mM Tris pH 7.5, 75 mM NaCl, 1% Triton X-100, 1 mM EDTA) supplemented with with cOmplete and PhosSTOP.

#### For western blotting

Cells (3.0 x 10^5^ in 6-well plates; 1.0 x 10^6^ in 100-mm dishes) were grown to confluence, washed twice with 1x PBS, and lysed in ice-cold lysis buffer (50 mM Tris-HCl pH 7.5, 150 mM NaCl, 1% Triton X-100, 1 mM EDTA) supplemented with with cOmplete and PhosSTOP. Lysates were processed as above, and boiled for 1 min before loading onto SDS-PAGE gels.

### GFP-trap and immunoprecipitation assays

For GFP-Trap assays, GFP-Trap Magnetic Agarose beads (ChromoTek) were washed with 1 mL of IP buffer. For each condition, 20 µL of beads were incubated with 500 µg protein extract for 2 h or overnight, at 4 °C under agitation. Beads were then washed twice with IP buffer, and bound proteins were eluted by adding 100 µL of 6x SDS loading buffer directly on the beads, followed by boiling.

For immunoprecipitation assays, Protein G agarose beads were washed with 1 mL TEN Buffer (Tris 50 mM pH 7.5, 1 mM EDTA, 100 mM NaCl), incubated with mouse anti-STARD3 (1STAR-2G5) in TEN Buffer (Di Mattia et al., 2018), and washed with IP buffer. The subsequent steps were performed as described for the GFP-Trap assays.

### Protein expression and purification

The recombinant fusion proteins consisting of the Avitag, His6 tag, MBP and TEV cleavage site either alone (thereafter named MBP) or fused to the MSP domains of VAP-A, VAP-B, MOSPD2 (WT and RD/LD mutant) were expressed in *E. coli* BL21 (DE3). Bacteria were cultivated in auto-inducible medium (Terrific Broth including trace elements; Formedium) supplemented with ampicillin (100 µg/mL) and containing 50 µM biotin (B4501 Merck) at 37°C until DO600 nm reached 0.6 and then placed at 20°C for 16 hours. Cells were pelleted and suspended in lysis buffer [50 mM Tris-HCl pH 7.5, 150 mM NaCl, 35 mM Imidazole, 1 mM DTT, 50 µM biotin, protease inhibitors tablets (cOmplete, Roche)]. Cells were lysed by a Cell Disruptor TS SERIES (Constant Systems Ltd), and the lysate was first centrifuged at 3,500 g for 15 minutes, then at 50,000 x g for 45 minutes, and filtered through a 0.45 µm membrane. Purification was performed on an ÄKTA Start chromatography system (GE Healthcare Life Sciences) using HisTrap HP 1 mL columns. Proteins were eluted with Elution buffer [50 mM Tris-HCl pH 7.5, 150 mM NaCl, 300 mM Imidazole, 1 mM DTT], and further purified by gel filtration (HiLoad 16/60 Superdex 200; GE) in GF Buffer [50 mM Tris-HCl pH 7.5, 150 mM NaCl, 2 mM DTT, 10 % Glycerol]. Proteins were concentrated with an Amicon Ultra-15 (30 kDa for MBP and 50 kDa for the other proteins) centrifugal filter unit (Merck). Protein concentration was determined by UV-spectroscopy.

### Native Holdup assays

For each MSP bait, 395 µL of streptavidin magnetic bead slurry (820 µL for the MBP control) (PureProteome Streptavidin Magnetic Beads, LSKMAGT10, Merck) in 1.5 mL tubes were washed twice with Holdup buffer. Beads were then incubated with 10 nmol of biotinylated MBP-MSP (40 nmol for MBP control) in a final volume of 750 µL (adjusted with Holdup buffer), for 2 h at 4°C under rotation. After saturation, beads were washed four times with 1 mL of buffer and resuspended to their initial slurry volume.

For native Holdup titration experiments, serial dilutions of MSP-saturated resin into control (MBP-saturated) resin were prepared in 0.5 mL tubes to obtain 100%, 50%, 25%, 12.5%, 6.25%, and 0% MSP-saturated resin, while keeping the total resin volume constant (200 µL slurry), and total molar amount of immobilized protein (MSP + MBP) constant. These conditions corresponded to apparent MSP bait concentrations of approximately 10, 5, 2.5, 1.25, 0.63, and 0 µM on the beads, respectively. After removal of the supernatant, resins were incubated with 100 µL of freshly prepared protein extracts (1 mg/mL supplemented with 1 mM DTT final) for 2 h at 4 °C under rotation and under an argon atmosphere. Unbound fractions were then separated using a magnetic rack and collected for Western blot analysis.

Western blot signals of unbound prey proteins were quantified using Fiji. In the native Holdup assay, binding is inferred from depletion of the prey in the unbound fraction. For each bait concentration, binding intensity (BI) was calculated as:

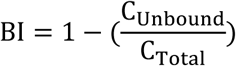

where C_Unbound_ corresponds to the prey signal intensity measured in the presence of the bait, and C_Total_ corresponds to the prey signal intensity measured with the MBP-saturated control resin (0% MSP), representing the total unbound prey.

Binding curves were generated in GraphPad Prism software using a specific binding with Hill slope nonlinear regression model to derive apparent binding parameters. For curve fitting, the mean value of technical replicates was used.

### SDS-PAGE and Western blot

SDS-PAGE and Western blot analysis were performed as previously described (Eichler et al., 2026) using the following antibodies: rabbit anti-GFP (1:2000; TP401, Torrey Pine Biolabs, RRID:AB_10013661), rabbit anti-MOSPD2 (1:1000; HPA003334, Sigma-Aldrich, RRID:AB_2146004), rabbit anti-mCherry (1:1000; ab167453, Abcam, RRID:AB_2571870), mouse anti-EEA1(1:1000; 610457, BD Biosciences, RRID:AB_397830), mouse anti-LAMP1 (1:1000; H4A3, DSHB, RRID:AB_2296838), mouse anti-LAMP2 (1:1000; H4B4, DSHB, RRID:AB_528129), mouse anti-VAP-A (1:1000; sc-293278, Santa Cruz Biotechnology, RRID:AB_2801294), anti-VAP-B [rabbit anti-VAP-B (Kabashi et al., 2013), rabbit anti-VAP-B (1:1000; HPA013144, Sigma-Aldrich, RRID:AB_1858717)], anti-STARD3 [rabbit polyclonal 605 (Alpy et al., 2001), mouse monoclonal 3G11 (Wilhelm et al., 2017)], rabbit anti-ORP1 (1:1000; ab131156, Abcam, RRID:AB_1155305), mouse anti-OSBP (1:1000; sc-365771, Santa Cruz Biotechnology, RRID:AB_10847232), rabbit anti-GAPDH (1:1000; G9545, Sigma-Aldrich, RRID:AB_796208) and mouse anti-actin (1:5000; ACT-2D7, Euromedex). To generate the anti-MOSPD2 rabbit polyclonal antibody #3452, a synthetic peptide corresponding to residues 1-17 (MAENH AQNKA KLISE TR C) with an added C-terminal cysteine was conjugated to ovalbumin and injected into New Zealand rabbits. The resulting serum was affinity purified and used for western blots.

For Coomassie staining, protein gels were stained with PageBlue Protein Staining Solution (ThermoFisher Scientific).

### Immunofluorescence (IF)

Cells (2.0 x 10^4^) were seeded onto 12-mm glass coverslips (No 1.5H, Marienfeld, 0117520) in 24-well plates, and IF experiments were performed 48 h after plating. For LysoTracker staining, cells were incubated with LysoTracker Red DND-99 (100 nM, ThermoFisher Scientific) in culture medium for 15 min at 37°C prior fixation. Cells were fixed in 4% paraformaldehyde in PBS for 15 minutes, washed twice in PBS, and permeabilized with 0.5 mg/mL filipin (F9765, Sigma-Aldrich) in PBS for 20 minutes (Eichler et al., 2025). After washes, and blocking with 1% bovine serum albumin in PBS (PBS-BSA), cells were incubated for 2h30 at room temperature with the primary antibody in PBS-BSA. Primary antibodies were: mouse anti-EEA1 (1:1000; 610457, BD Biosciences, RRID:AB_397830), mouse anti-LAMP1 H4A3 (1:100; DSHB, RRID:AB_2296838). For BMP labelling using mouse anti-BMP 6C4 (10 µg/mL; kind gift from Toshihide Kobayashi) and STARD3 labelling using rabbit polyclonal 1611 (Alpy et al., 2001), cells were permeabilized with 50 µg/mL digitonin for 10 minutes instead of filipin. Cells were washed twice in PBS and incubated for 30 minutes with the secondary antibody (1:1000; AlexaFluor 488 (RRID: AB_2535792 and AB_141607), AlexaFluor 555 (RRID: AB_2762848 and AB_162543), AlexaFluor 647 (RRID: AB_2536183 and AB_162542) from ThermoFisher Scientific), and Hœchst-33258 (1:10 000; B2261, Sigma-Aldrich) to stain nuclei. After two washes with PBS, coverslips were mounted in ProLong Gold (Invitrogen).

Images were acquired using a spinning disk CSU-X1 (Nikon, 100x NA 1.4 or 63x NA 1.4), and for filipin imaging with a Leica SP8 UV inverted confocal microscope (63X, NA 1.4). Laser power, z-stack parameters, and exposure settings were kept constant within each experiment.

### Live-cell imaging

For live-cell imaging, 9 x 10^5^ mClover3-MOSPD2 cells were seeded in 35 mm glass-bottom dishes (MatTek). Forty-eight hours later, cells were incubated for 15 min in phenol red-free culture medium supplemented with LysoTracker (100 nM) to label LE/Lys. Imaging was performed on a laser-scanning microscope equipped with Airyscan (Zeiss LSM 980) with a 63x oil immersion objective (NA 1.4). Z-stacks were acquired with 0.15 µm step size (41 slices).

Airyscan SR joint deconvolution was applied using the microscope software, and images were subsequently processed using Fiji (Z-projection of 6 consecutive slices).

### Quantification of endosome number and morphology

For image processing, a z-stack projection (max intensity) was generated in ImageJ/Fiji (Schindelin et al., 2012). Endosomes were segmented using a custom StarDist model (Schmidt et al., 2018). First, images of endosomes stained with anti-STARD3, anti-LAMP1 antibodies, and with LysoTracker were cropped, and 127 images were selected for training. These images were manually annotated using QuPath (Bankhead et al., 2017). A StarDist model was then trained online on a Google Colab notebook created by ZeroCostDL4Mic platform (von Chamier et al., 2021). This custom model was used locally using Python to perform instance segmentation. The segmented images were processed in CellProfiler (McQuin et al., 2018) where cells were manually segmented by rescaling the intensity of the original images. Multiple parameters (object intensity, object neighbors and object size/shape) were analyzed for the identified endosomes. The resulting data were then processed using Spyder 4.1 (Python 3.7) and GraphPad Prism.

### Quantification of LE/Lys spatial distribution

LE/Lys spatial distribution was quantified using CellProfiler (version 4.2.8). Nuclei were automatically segmented from the Hoechst channel (IdentifyPrimaryObjects), and cell boundaries were manually defined. Measurements were restricted to the cytoplasm between the nuclear periphery and the cell edge. LE/Lys were segmented as described above, then reduced to single-pixel objects (ExpandOrShrinkObjects) and converted into a binary image (ConvertObjectsToImage), to avoid size or intensity biases. Radial distributions were computed using the MeasureRadialDistribution module. Radial distances were normalized from the nuclear periphery (distance = 0) to the cell boundary (distance = 1). Each cell was subdivided into two concentric radial bins of equal normalized thickness, and the fraction LE/Lys in each bin was measured (FracAtD). Because the input image was binary, these fractional intensity values directly reflected the fraction of LE/Lys located within each radial region.

### Quantification of LE/Lys tubules undergoing fission

The proportion of LE/Lys tubules undergoing fission was quantified as previously described (Boutry et al., 2023). U2OS cells were silenced for MOSPD2 using siRNA and transfected with Lamp1-mCherry plasmid to label LE/Lys membranes. Cells were plated on Lab-Tek chambers (Nunc) with #1.5 borosilicate glass bottoms. To induce tubule formation, cells were starved for 10 h in HBSS (Hanks’ Balanced Salt Solution, Gibco) and then imaged using a Leica SP8 laser scanning confocal microscope equipped with a Lightning deconvolution module, a 63× glycerol immersion objective lens (NA 1.3), at 37°C and 5% CO_2_. Images were acquired every 1.5 s for 2 min. To account for the increased number of LE/Lys vesicles in MOSPD2-depleted cells, we quantified the proportion of tubules undergoing fission. To standardize measurements, at the beginning of each movie, 10 random LE/Lys tubules were selected, and their fate (fission vs. no fission) was monitored. LE/Lys tubule fission events were defined as events showing a clear division of a Lamp1-mCherry-positive tubule emerging from a Lamp1-mCherry-positive organelle.

### Quantification of LE/Lys fusion

Cells (2.0 x 10^5^) were seeded on glass coverslips in 24-well plates. The following day, cells were incubated with Alexa Fluor 647-labeled dextran (2 µg/mL; D22914, ThermoFisher Scientific) for 6 h, washed, and incubated overnight in standard culture medium; this allowed the loading of LE/Lys with Alexa Fluor 647-dextran. The next day, cells were incubated with Oregon Green 488-labeled dextran (0.2 mg/mL; D7170, ThermoFisher Scientific) for 30 min, washed with standard culture medium, and fixed at different chase time points (0, 20, 40 min). Coverslips were mounted in ProLong Gold (Invitrogen). Images were acquired on a spinning disk CSU-X1 (Nikon, 100x NA 1.4), using identical acquisition settings across conditions (laser power, z-slices, and exposure).

Alexa Fluor 647– and Oregon Green 488-labeled vesicles were segmented using a custom Stardist model, as described above. Image analysis was subsequently performed using CellProfiler. Vesicles that had undergone fusion were defined as new object generated by colocalization of Alexa Fluor 647– and Oregon Green 488-positive vesicles. The proportion of fused vesicles was calculated as the number of colocalizing vesicles divided by the vesicle count (sum of both channels minus the colocalizing population).

### Fluorescence Recovery After Photobleaching (FRAP)

COS7 cells were plated on 35 mm glass-bottom dishes (MatTek), transfected with plasmids encoding GFP-MOSPD2, GFP-VAPA, GFP-VAPB with mCherry-STARD3, and allowed to grow for 48 hours. Experiments were performed using a spinning disk CSU-X1 (Nikon, 100x oil objective, NA 1.4). Regions of interest were photobleached with the 405 nm laser line at 15% power and 5 repetitions. Fluorescence recovery was monitored every 250 ms for 5 seconds, then every second for 15 seconds after photobleaching.

For image processing, tracking of the intensity from the ROI was performed on ImageJ using an in-house script. Non-bleached endosomes were also tracked to measure natural photobleaching of the dye. Data were then analyzed using Spyder 4.1 (Python 3.7). Each ROI was normalized by dividing the raw intensity of each frame by the intensity of the first frame before bleaching.

### GFP-D4-based quantification of cholesterol

Cells (2.0 x 10^6^) were seeded in T75 flasks and cultured for 48 hours. Then, 3.0 x 10^5^ cells were plated onto glass coverslips in 24-well plates, and grown for another 48 hours. Cells were then incubated with 100 nM LysoTracker Red DND-99 for 15 minutes prior to fixation. Cells were fixed in 4% PFA (PBS 1X) and permeabilized with a brief bath (1 second) in liquid nitrogen. After 30 min blocking in 1% BSA (PBS 1X), cells were incubated with GFP-D4 (Wilhelm et al., 2019) or mEGFP-D4H (Eichler et al., 2025) (100 μg/mL in blocking solution) for 2 hours at room temperature. Cells were washed and fixed again in 4% PFA (PBS 1X) for 10 mins. After two washes, coverslips were mounted in ProLong Gold (Invitrogen). Images were acquired on a spinning disk CSU-X1 (Nikon, 100x, NA 1.4), using consistent settings for laser power, z-slices, and exposure.

Endosomes were segmented using a custom StarDist model. GFP-D4 intensity was quantified in LysoTracker-positive objects using CellProfiler. Data were normalized to the mean intensity of WT cells using Spyder 4.1 (Python 3.7).

### Filipin-based quantification of cholesterol

Cells (2.0 x 10^6^) were seeded in T75 flasks and cultured for 48 hours. Then, 3.0 x 10^5^ cells were plated on glass coverslips in 24-well plates, and grown for another 48 hours. Cells were fixed in 4% PFA in PBS 1X and permeabilized with filipin. LE/Lys were labeled using anti-LAMP1 antibodies. Coverslips were mounted in ProLong Gold (Invitrogen). Confocal images were acquired on a Leica SP8-UV inverted microscope (63x, NA 1.4), using identical settings for laser power, z-slices, and exposure across all experiments.

LE/Lys were segmented using a custom StarDist model. Images were then processed on CellProfiler. Cells were manually segmented using a rescaled intensity of the filipin channel. Filipin intensity within LAMP1-positive structures was measured. Data were normalized to the mean intensity of WT or siCtrl conditions using Spyder 4.1 (Python 3.7).

### Transmission electron microscopy

Electron microscopy was performed as previously described (Alpy et al., 2013; Zouiouich et al., 2022). Cells grown on carbon-coated sapphire disks were cryoprotected with DMEM containing 10% FCS and frozen at high pressure (HPM 10 Abra Fluid AG). Samples were then freeze-substituted and embedded in lowicryl HM20. Thin sections were collected on formvar-/carbon-coated nickel slot grids and stained with uranyl acetate and lead citrate. Imaging was performed with a transmission electron microscope (FEI TECNAI G2-20 Twin) coupled to a TemCam-F216 camera (TVIPS).

### Statistical analyses

Statistical analyses were performed using one-way ANOVA or Student’s t-test (Prism, GraphPad). Multiple comparisons were conducted using Tukey’s or Dunnett’s test, as appropriate. P values <0.05, <0.01, <0.001 and <0.0001 are identified with *, **, *** and ****, respectively. “ns” indicates P ≥ 0.05.

## Figure legends

**Figure S1:**
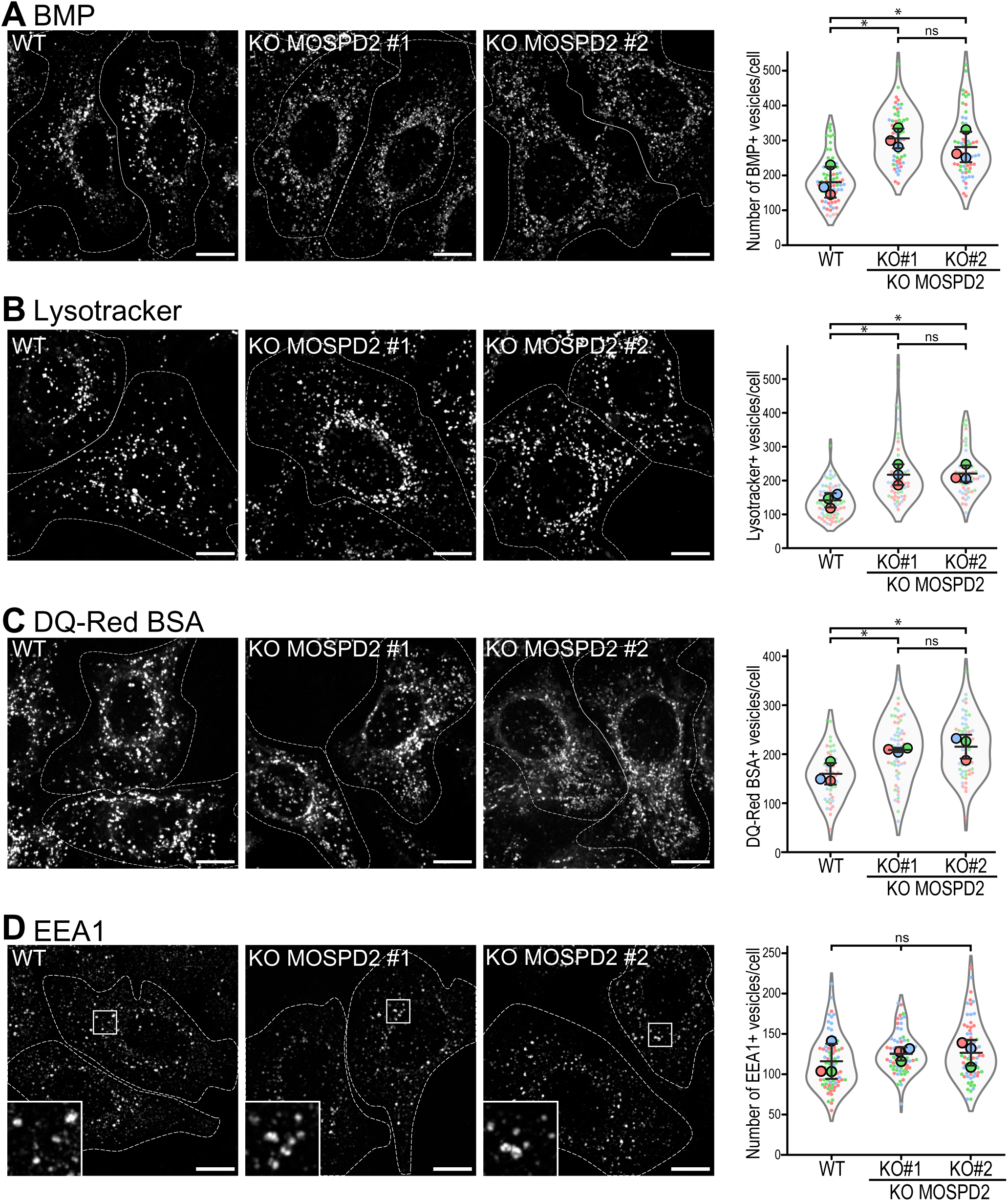
Characterization of LE/Lys in MOSPD2-deficient cells. Representative confocal images of parental HeLa cells (WT) and MOSPD2-deficient clones (MOSPD2 KO#1 and MOSPD2 KO#2) labeled with anti-BMP antibodies (A; gray), Lysotracker (B; gray), DQ-Red BSA (C; gray), and EEA1 (D; gray). Scale bars: 10 µm. The contour of each cell is delimited by a white dotted line. Right: quantification of the numbers of BMP– (A), Lysotracker– (B; gray), DQ-Red BSA (C; gray), and EEA1-positive vesicles under the conditions shown on the Left. Data are displayed as Superplots showing the mean per cell (small dots) and per independent experiment (large dots). Independent experiments (n = 3) are color-coded. Means and error bars (SD) are shown as black bars. Data were collected from A: 79 (WT), 60 (KO#1), and 63 (KO#2) cells; B: 76 (WT), 59 (KO#1), and 58 (KO#2) cells; C: 60 (WT), 56 (KO#1), and 53 (KO#2) cells D: 80 (WT), 76 (KO#1), and 73 (KO#2) cells. One-way ANOVA with Tukey’s multiple comparisons test (ns, not significant; *, P < 0.05; n = 3 independent experiments).

**Figure S2:**
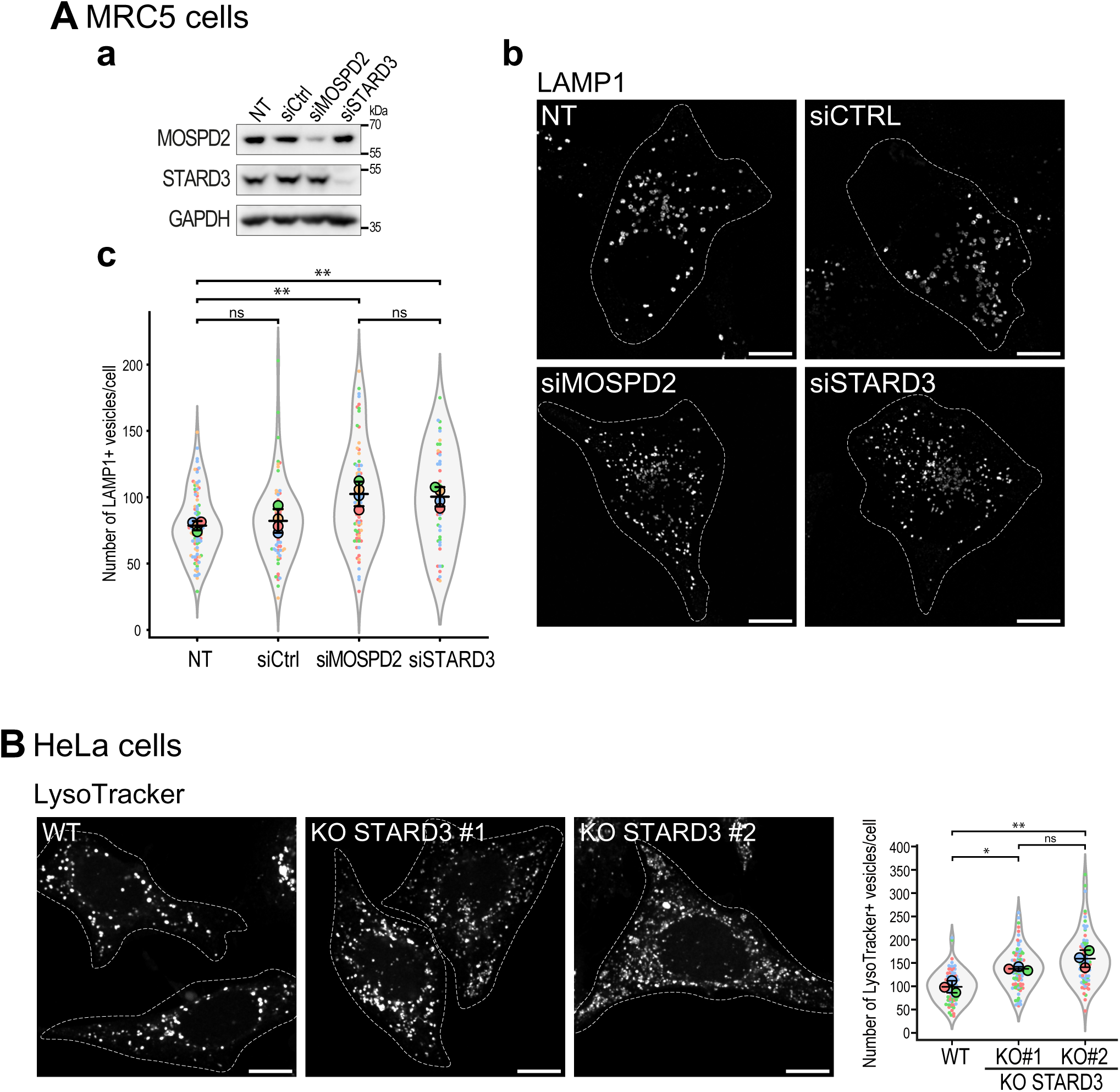
Characterization of LE/Lys in MOSPD2– or STARD3-deficient MRC5 and HeLa cells. A: a: Western blot analysis of MOSPD2, STARD3 and GAPDH expression levels in control MRC5 cells (NT and siCtrl) and in cells treated with siRNAs targeting MOSPD2, and STARD3. b: Representative confocal images of LAMP1 labeling (grey) in control MRC5 cells (NT, not transfected; and siCtrl) and in cells tranfected with siRNAs targeting MOSPD2 and STARD3. The contour of each cell is delimited by a white dotted line. Scale bars: 10 µm. c: quantification of the number of LAMP1-positive vesicles under the conditions shown in b. Data are displayed as Superplots showing the mean per cell (small dots) and per independent experiment (large dots). Independent experiments (n = 4) are color-coded. Means and error bars (SD) are shown as black bars. Data were collected from 95 (NT), 74 (siCtrl), 76 (siMOSPD2), and 49 (siSTARD3). One-way ANOVA with Tukey’s multiple comparisons test (ns, not significant; **, P < 0.01; n = 4 independent experiments). B: Left: Representative confocal images of parental HeLa cells (WT) and STARD3-deficient clones (KO#1 and KO#2) labeled with Lysotracker (gray). Scale bars: 10 µm. The contour of each cell is delimited by a white dotted line. Right: quantification of the numbers of Lysotracker-positive vesicles under conditions shown on the Left. Data are displayed as Superplots showing the mean per cell (small dots) and per independent experiment (large dots). Independent experiments (n = 3) are color-coded. Means and error bars (SD) are shown as black bars. Data were collected from 81 (WT), 95 (KO#1), and 86 (KO#2) cells. One-way ANOVA with Tukey’s multiple comparisons test (ns, not significant; *, P < 0.05; **, P < 0.01; n = 3 independent experiments).

**Figure S3:**
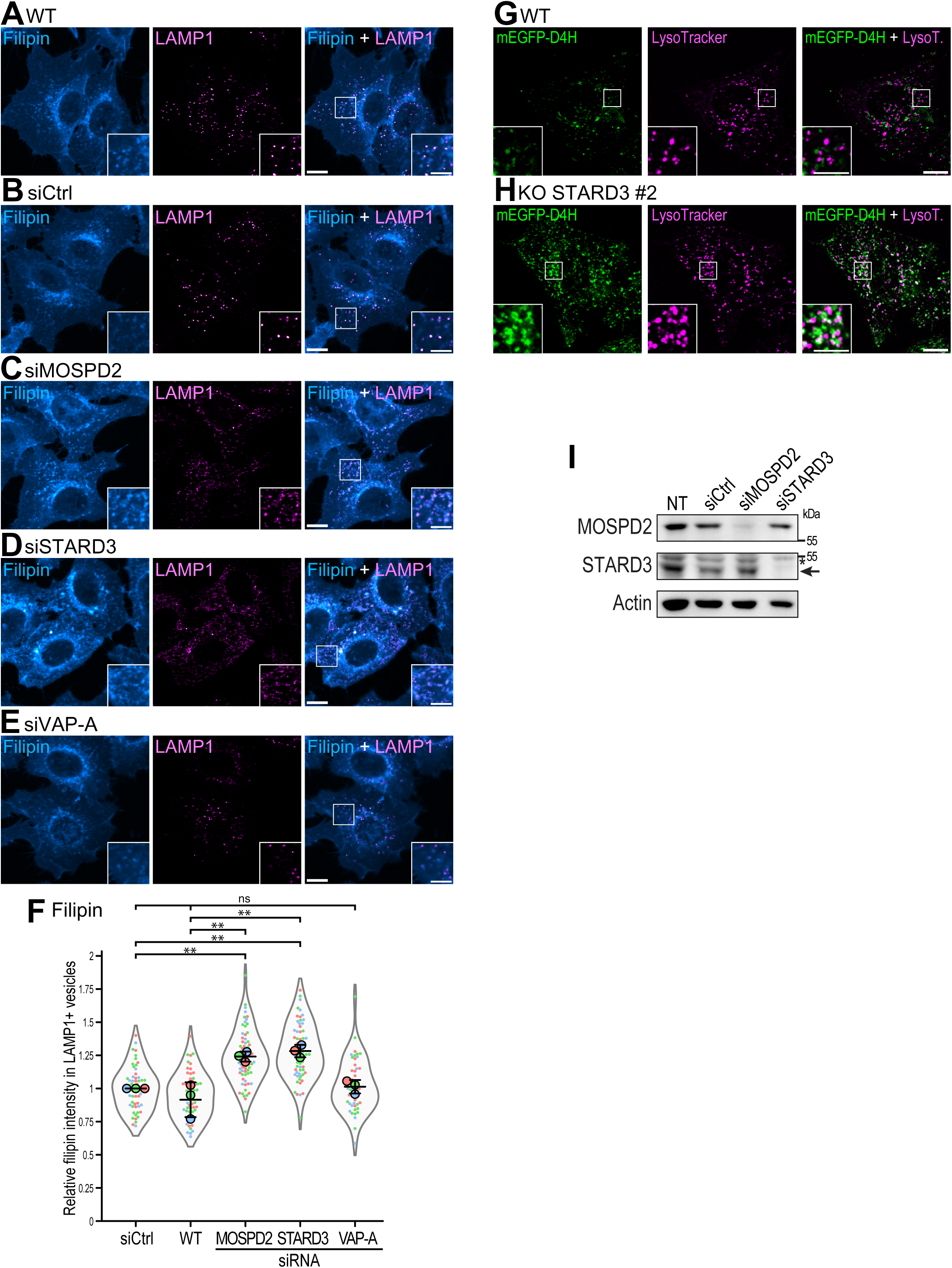
MOSPD2– and STARD3-deficient cells harbor increased cholesterol levels in LE/Lys. A-E: HeLa cells either untransfected (A) or transfected with control siRNAs (B) or siRNAs targeting MOSPD2 (C), STARD3 (D) or VAP-A (E) were labeled with anti-LAMP1 antibodies (magenta Hot) and the fluorescent cholesterol probe filipin (Cyan Hot). Higher magnifications images of the area outlined in white are shown on the right. Scale bars: 10 µm; inset scale bars: 5 µm. F: Quantification of relative filipin fluorescence intensity in LAMP1-positive vesicles under the conditions shown in A-E. Data are presented as Superplots, displaying the mean number of LE/Lys per cell (small dots) and per independent experiment (large dots). Independent experiments (n = 3) are color-coded. Means and SD are indicated as black bars. Data were collected from 53 (siCtrl), 59 (WT), 69 (siMOSPD2), 64 (siSTARD3), 49 (siVAPA) cells. One-way ANOVA with Tukey’s multiple comparisons test (ns, not significant; **, P < 0.01; n = 3 independent experiments). G-H: WT (G) or STARD3 KO (H) HeLa cells were labeled with the fluorescent cholesterol probe GFP-D4 (green), and Lysotracker (magenta). Higher magnifications images of the area outlined in white are shown. Scale bars: 10 µm; inset scale bars: 5 µm. I: Western blot analysis of MOSPD2, STARD3, and Actin protein levels in HeLa cells stably expressing the mEGFP-D4H probe, either untransfected (NT) or transfected with control siRNAs (siCtrl) or siRNAs targeting MOSPD2 (siMOSPD2) or STARD3 (siSTARD3).

**Figure S4:**
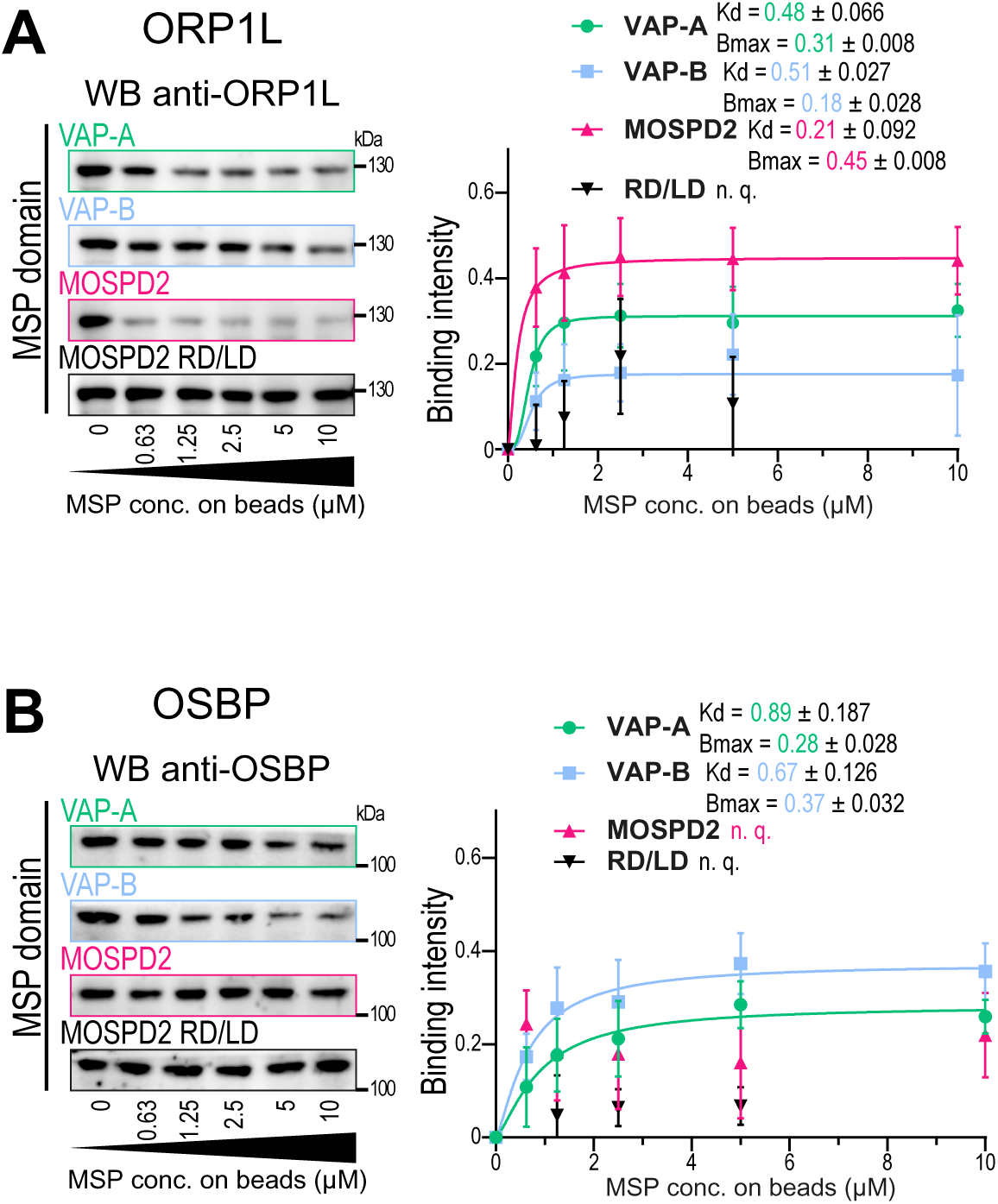
Quantification of the interactions between ORP1L, OSBP, and the MSP domains of VAP-A, VAP-B and MOSPD2. Native Holdup experiments quantifying the interaction between the recombinant MSP domains of VAP-A, VAP-B, MOSPD2, and MOSPD2 RD/LD and endogenous ORP1L (A) and OSBP (B). Left: western blot analysis of the unbound prey protein in HCC1954 protein extracts after incubation with increasing amounts of the recombinant MSP domains. Right: Binding intensity between the MSP domains and the prey proteins. Binding curves were fitted using a Hill equation (mean ± SEM from 4-5 technical replicates for ORP1L and 3 technical replicates for OSBP), and apparent affinities (*K*_app_) and maximal binding intensities (*B*_max_) were calculated (± SD).

## Acknowledgments

We thank Alastair McEwen for his critical reading of the manuscript. We thank the members of the Molecular and Cellular Biology of Breast Cancer team (IGBMC) and the members of Toshihide Kobayashi’s team for helpful advice and discussions. We are grateful to the members of the IGBMC Imaging Center (ICI), especially Elvire Guiot and Erwan Grandgirard. We acknowledge the IGBMC imaging center, member of the national infrastructure France-BioImaging supported by the French National Research Agency (ANR-10-INBS-04). We thank Chadia Nahy from the IGBMC Electron Microscopy Facility and Rachel Mellwig, Paolo Ronchi and Yannick Schwab from the Electron Microscopy Core Facility at the EMBL Heidelberg for their help with electron microscopy. We thank the IGBMC cell culture facility (Amelie Freismuth), the Flow Cytometry facility (Claudine Ebel and Muriel Philipps), the Molecular Biology and Virus Service (Paola Rossolillo and Nicole Jung), and the Mediaprep facility (Denis Fumagalli).

C. Knorr received fellowships from L’Alsace contre le Cancer (https://alsacecontrecancer.com/) and from SEVE Sein et Vie. M. Zouiouich received fellowships from ITMO Cancer AVIESAN (Alliance Nationale pour les Sciences de la Vie et de la Santé/National Alliance for Life Sciences and Health) within the framework of the Cancer Plan (https://itcancer.aviesan.fr/) and from Fondation ARC pour la recherche sur le cancer (ARCDOC42022010004428). S. Huver received a fellowship from the Ligue Nationale contre le Cancer. J. Eichler received fellowships from EUR IMCBio funds and the Ligue Nationale contre le Cancer.

This work was supported by grants from the Agence Nationale de la Recherche ANR (grant ANR-25-CE44-6381 and ANR-21-CE13-0014; https://anr.fr/), and from the Ligue Contre le Cancer (Conférence de Coordination Interrégionale du Grand Est; https://www.ligue-cancer.net).

This work of the Interdisciplinary Thematic Institute IMCBio, as part of the ITI 2021-2028 program of the University of Strasbourg (http://www.unistra.fr), CNRS (http://www.cnrs.fr/) and Inserm (http://www.inserm.fr/), was supported by IdEx Unistra (ANR-10-IDEX-0002), and by SFRI-STRAT’US project (ANR 20-SFRI-0012) and EUR IMCBio (ANR-17-EURE-0023) under the framework of the French Investments for the Future Program.

